# Pex24 and Pex32 tether peroxisomes to the ER for organelle biogenesis, positioning and segregation

**DOI:** 10.1101/2020.03.05.977884

**Authors:** Fei Wu, Rinse de Boer, Arjen M. Krikken, Arman Akşit, Nicola Bordin, Damien P. Devos, Ida J. van der Klei

## Abstract

We analyzed all four Pex23 family proteins of the yeast *Hansenula polymorpha*, which localize to the ER. Of these Pex24 and Pex32, but not Pex23 and Pex29, accumulate at peroxisome-ER contacts, where they are important for normal peroxisome biogenesis and proliferation and contribute to organelle positioning and segregation.

Upon deletion of *PEX24* and *PEX32* - and to a lesser extent of *PEX23* and *PEX29* - peroxisome-ER contacts are disrupted, concomitant with peroxisomal defects. These defects are suppressed upon introduction of an artificial peroxisome-ER tether.

Accumulation of Pex32 at peroxisomes-ER contacts is lost in the absence of the peroxisomal membrane protein Pex11. At the same time peroxisome-ER contacts are disrupted, indicating that Pex11 contributes to Pex32-dependent peroxisome-ER contact formation.

Summarizing, our data indicate that *H. polymorpha* Pex24 and Pex32 are tethers at peroxisome-ER contacts that are important for normal peroxisome biogenesis and dynamics.

**Summary:** Two *Hansenula polymorpha* ER proteins, Pex24 and Pex32, are tethers at peroxisome-ER contacts and function together with the peroxisomal protein Pex11. Their absence disturbs these contacts leading to multiple peroxisomal defects, which can be restored by an artificial tether.

## Introduction

Peroxins are defined as proteins that play a role in peroxisome biogenesis, including peroxisomal matrix protein import, membrane biogenesis and organelle proliferation (Distel et al., 1996). Most peroxins are peroxisomal or cytosolic proteins, which transiently can be recruited to the organelle. Recent studies in bakers’ yeast showed that a family of peroxins, called the Pex23 protein family (Kiel et al., 2006), localize to the endoplasmic reticulum (ER) (David et al., 2013; Joshi et al., 2016; Mast et al., 2016). The function of these peroxins is still poorly understood and is the subject of this study.

Proteins of the Pex23 family are characterized by a DysF domain. The DysF domain was first identified in human dysferlin. Dysferlin is important for fusion of vesicles with the sarcolemma at the site of muscle injury (Bansal and Campbell, 2004; North et al., 2011; Bansal et al., 2003). Dysferlin contains multiple C2 domains, which play a direct role in the above membrane repair process, however the function of the DysF domain is still obscure.

*Yarrowia lipolytica* Pex23 was the first DysF domain containing peroxin that was identified (Brown et al., 2000). All DysF domain containing yeast peroxins belong to the Pex23 protein family. The number of Pex23 family members varies in different yeast species and their nomenclature is confusing (see Fig. 1A). *Hansenula polymorpha* and *Pichia pastoris* contain four members, but *Saccharomyces cerevisiae* has five and *Y. lipolytica* has three. Mutants lacking one of these peroxins show diverse peroxisomal phenotypes ranging from a partial matrix protein import defect to enhanced or decreased peroxisome numbers (Brown et al., 2000; Tam and Rachubinski, 2002; Vizeacoumar et al., 2003, 2004; Yan et al., 2008).

**Figure 1.**
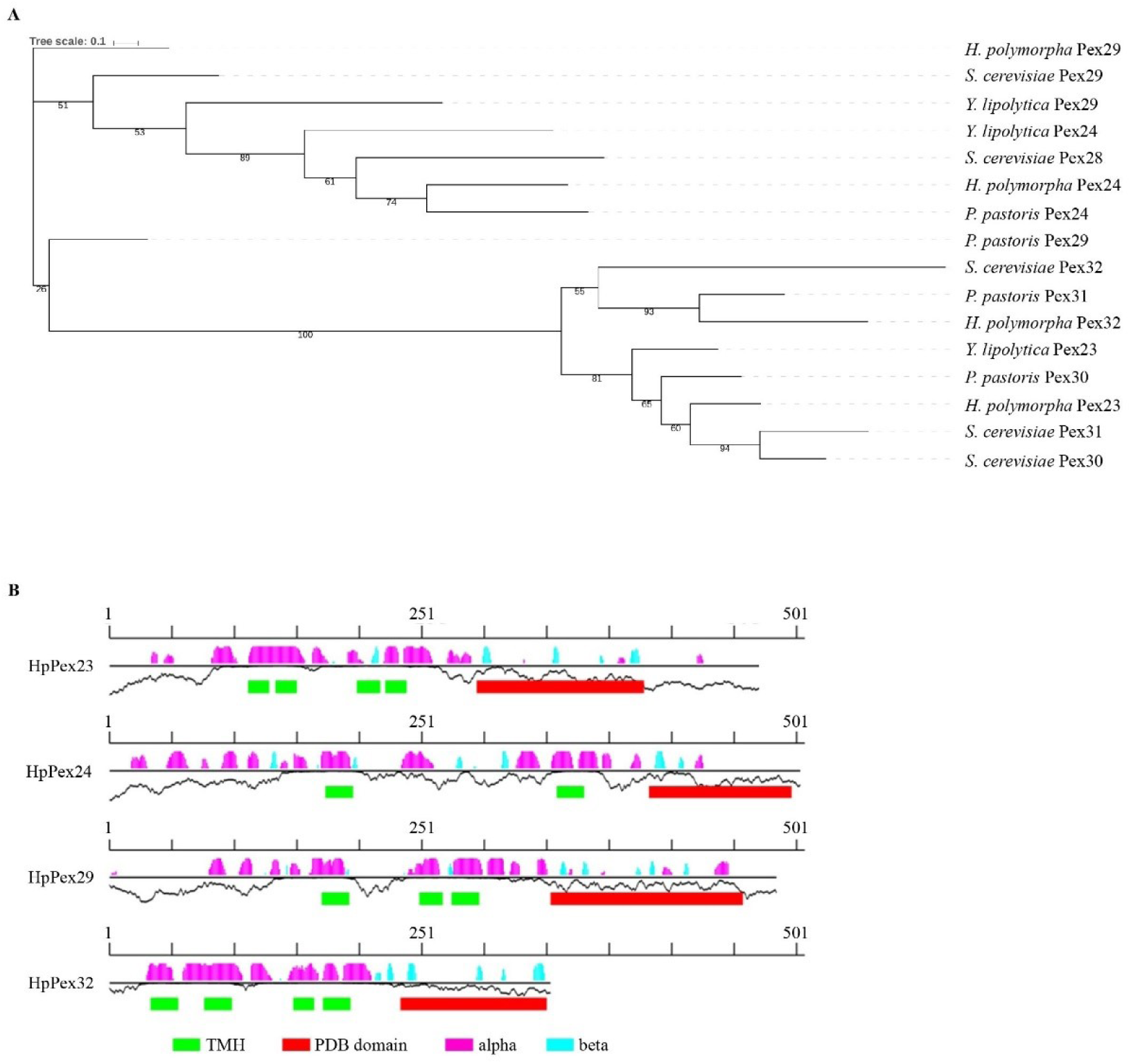
Yeast Pex23 family proteins. **(A)** Protein phylogeny. Protein sequences from *S. cerevisiae, H. polymorpha, P. pastoris* and *Y. lipolytica* were retrieved from NCBI-protein. Phylogenetic tree: numbers represent the bootstraps values, while branch length represents the amino acidic substitution rates. (**B)** Secondary structure features of *H. polymorpha* Pex23 proteins obtained with Foundation (Bordin et al., 2018). The black horizontal lines represent the protein sequence. The predicted β-strands and α-helices are depicted by bars above each line in cyan and magenta, with the height of the bars representing the confidence of the prediction. Transmembrane helices (TMH) predictions are depicted as green boxes underneath the secondary structure prediction. The Protein Data Bank (PDB) domain represents the DysF domain.

Initially, proteins of the Pex23 family were proposed to localize to peroxisomes (Brown et al., 2000; Tam and Rachubinski, 2002; Vizeacoumar et al., 2003, 2004; Yan et al., 2008). However, later studies indicated that *S. cerevisiae* Pex23 family proteins are ER proteins and form complexes with the ER resident reticulons Rtn1/Rtn2 and Yop1 (David et al., 2013; Mast et al., 2016). *S. cerevisiae* Pex30 and its paralog Pex31 have been implicated in the formation of ER-peroxisome contact sites and are suggested to regulate *de novo* peroxisome formation from the ER (David et al., 2013; Mast et al., 2016). *S. cerevisiae* Inp1 also has been implicated in the formation of peroxisome-ER contacts, but serves a different function, namely in peroxisome retention during yeast budding (Knoblach et al., 2013).

ScPex30 and ScPex31 contain a reticulon-like domain and have membrane shaping properties (Joshi et al., 2016). Regions at the ER where Pex30 accumulates are important for de novo peroxisome formation, but also play a role in lipid droplet biogenesis (Joshi et al., 2018; Wang et al., 2018; Lv et al., 2019).

So far, *S. cerevisiae* Pex29, Pex30 and Pex31 have been extensively studied. However, our knowledge on other members of the *S. cerevisiae* Pex23 protein family as well as on these proteins from other yeast species is still relatively scarce.

Here, we systematically studied all four Pex23 family members from the yeast *Hansenula polymorpha*. Our results indicate that these proteins localize to the ER and accumulate at specific membrane contact sites, including peroxisome-ER contacts and nucleus vacuole junctions (NVJs). Deletion of *PEX24* or *PEX32* results in major aberrations in peroxisome biology. These defects are accompanied by enhanced distances between peroxisomal and ER membranes. Introduction of an artificial peroxisome-ER tether suppresses these peroxisomal phenotypes, suggesting that Pex24 and Pex32 function as tethers at contact sites.

The absence of the peroxisomal membrane protein Pex11 also disrupts ER-peroxisome contacts, accompanied by reduced accumulation of Pex32 at these contacts. These results are consistent with the view that Pex11 functions together with Pex23 family proteins to associate peroxisomes to the ER.

Summarizing, our data show that Pex24 and Pex32 play key roles in the association of peroxisomes to the ER. The loss of these associations affects multiple organelle properties, including peroxisome biogenesis, positioning and segregation.

## RESULTS

### Protein sequence and structure prediction

Construction of a phylogenetic tree of Pex23 family members of four different yeast species indicated that two subfamilies (the Pex23 and Pex24 subfamilies) can be distinguished (Fig. 1A). All *H. polymorpha* members contain a DysF domain at the C-terminus. HpPex32 is much shorter than the others, which is mostly due to the lack of an unstructured fragment in this protein (Fig 1B).

HpPex23 ends with a KKKE stretch of residues, similar to the KKXX found in *S. cerevisiae* Pex30 (David et al., 2013). *H. polymorpha* Pex24 ends with KKR. These C-termini may represent di-lysine motifs, which are recognized by coatomer subunits and important for retrograde transport to the ER (Ma and Goldberg, 2013). The C-termini of HpPex29 and HpPex32 do not contain di-lysine motifs.

Secondary structure prediction indicated that all four sequences contain between two to four transmembrane helices and a C-terminal domain dominated by beta-sheets (Fig. 1B). It has been previously argued that a reticulon-like domain was observed in this family of proteins, particularly in ScPex30 and ScPex31 (Joshi et al., 2016). Indeed, a similar hit can be found on HpPex23 using HHpred on the Pfam-A database. This detection extends from residue 100 to 233 of HpPex23. However, this detection has an E-value of 2 with a probability of 92.38, making it a borderline detection. Similar borderline hits are detected in HpPex24, HpPex29 and HpPex32. A Trp residue is also present at the N terminus of this potential domain and aligns with the classical Trp conserved residue of other Pex reticulon-like domains.

### All *H. polymorpha* Pex23 family members localize to the ER

The localization of the four *H. polymorpha* Pex23 family proteins was determined by fluorescence microscopy (FM) using strains producing C-terminal GFP tagged proteins under control of their endogenous promoters (Fig. 2A,B). Cells were grown in media containing glucose (peroxisome repressing conditions). At these conditions the cells generally contain a single small peroxisome, associated to the ER (Wu et al., 2018). As shown in Fig. 2B, all four proteins co-localized with the ER marker BiP-mCherry-HDEL, predominantly at the cortical ER. Frequently, a patch of Pex23-GFP was observed at the nuclear envelope as well (Fig. 2A, B). In Pex24-GFP or Pex32-GFP producing strains generally one fluorescent spot was detected in each cell, which invariably localized close to the single peroxisome marked with Pex14-mKate2. More spots were present in cells of Pex23-GFP and Pex29-GFP producing strains, but one of these invariably was present in the vicinity of the Pex14-mKate2 spot (Fig. 2A).

**Figure 2.**
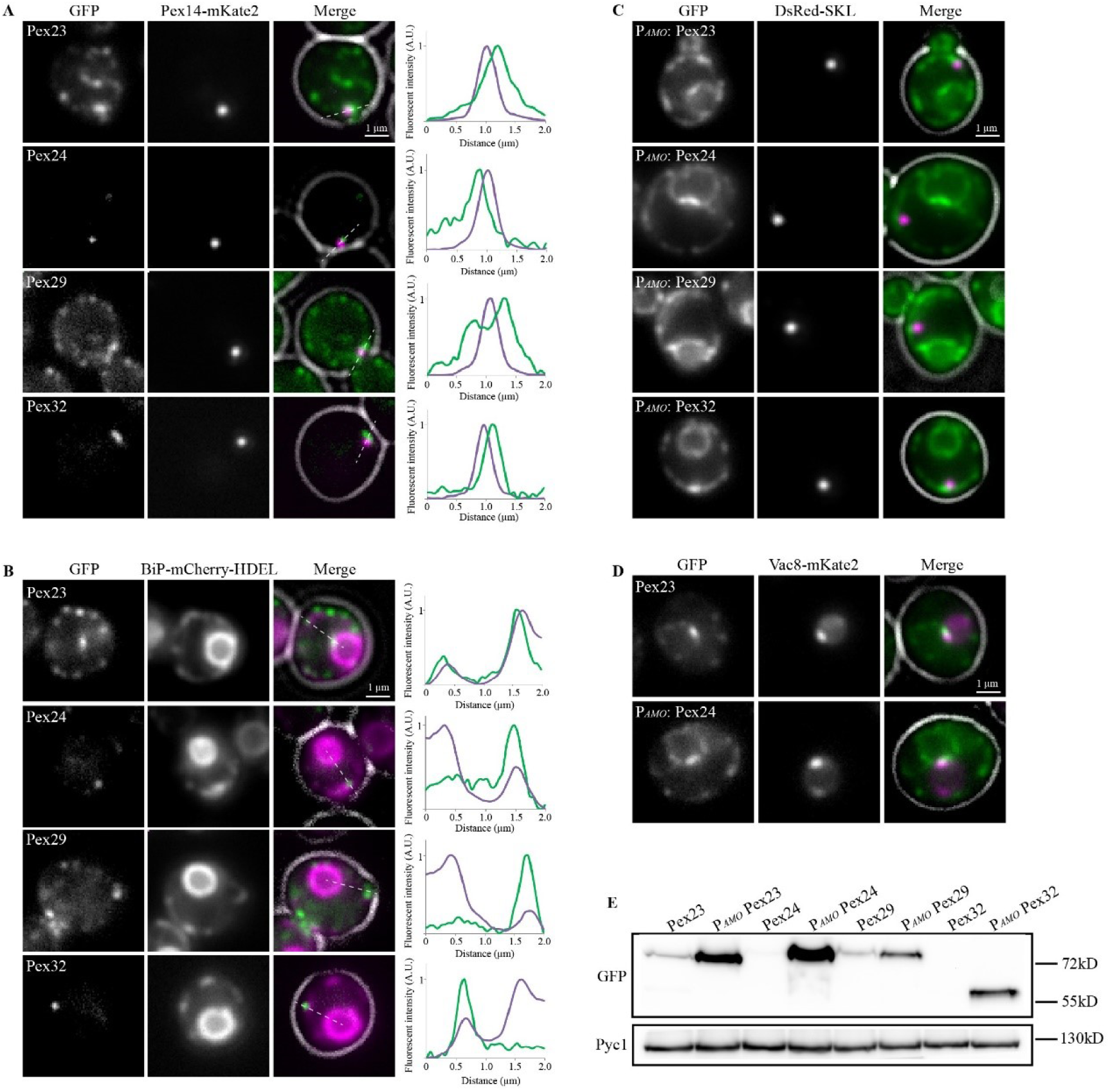
*H. polymorpha* Pex23 family proteins localize to the ER. FM images of glucose-grown *H. polymorpha* cells producing the indicated GFP fusion proteins under control of their endogenous promoters together with the peroxisomal marker Pex14-mKate2 **(A)** or the ER marker BiP-mCherry-HDEL **(B). (A**,**B)** The merged images show the cell contours in white. Graphs show relative fluorescence intensity along the dotted lines. **(C)** FM images of glucose/methylamine-grown *H. polymorpha* WT cells producing the peroxisomal marker DsRed-SKL and the indicated GFP fusion proteins under control of the amine oxidase promoter (P_*AMO*_). **(D)** Co-localization of Vac8-mKate2 with Pex23-GFP produced under control of the endogenous promoter or Pex24-GFP expressed under control of the P_*AMO*_. **(E)** Western Blot analysis of the indicated strains. Cells were grown for 4 hours on glucose. Strains producing the GFP fusion proteins under control of the P_*AMO*_ were grown in media containing methylamine as nitrogen source. Equal amounts of cellular lysates were loaded per lane. Blots were decorated with a-GFP or α-Pyruvate carboxylase 1 (Pyc1) antibodies. Pyc1 was used as a loading control.

Upon overproduction, all four HpPex23 proteins showed a typical cortical ER/nuclear envelope pattern, supporting that they represent genuine ER proteins. FM analysis revealed that the overproduced proteins were not evenly distributed over the ER, but present in spots and patches. In all strains one cortical patch localized in the vicinity of the peroxisome (here marked with DsRed-SKL) (Fig. 2C). Relatively large patches of GFP fluorescence were frequently observed at the nuclear envelope in cells overproducing Pex23-GFP or Pex24-GFP. Co-localization studies with the nucleus-vacuole junction (NVJ) protein Vac8 indicated that these patches represent NVJs (Fig. 2D). Pex23-GFP also accumulated at NVJs when produced under control of its own promoter.

Western blot analysis showed that the levels of all four GFP fusion proteins were very low when produced under control of their endogenous promoters. In fact, Pex24-GFP and Pex32-GFP were below the limit of detection, whereas faint bands were detected on blots of Pex23-GFP and Pex29-GFP producing cells. Upon overproduction all four GFP-fusion proteins were readily detected (Fig. 2E).

Our data support observations in *S. cerevisiae*, where Pex23 family proteins localize to the ER, including at regions where peroxisomes and ER are in close vicinity (David et al., 2013; Mast et al., 2016). The presence of a portion of HpPex23 and overproduced HpPex24 at NVJs suggest that Pex23 family proteins are also components of other membrane contacts.

### Pex24 and Pex32 are important for peroxisome biogenesis and proliferation

To study the role of the Pex23 family proteins we constructed four *H. polymorpha* deletion strains, *pex23, pex24, pex29* and *pex32*.

First, we analyzed whether Pex23 family proteins are important for peroxisomal matrix protein import using glucose-grown cells producing the matrix marker GFP-SKL and wide field fluorescence microscopy (FM). GFP-SKL mislocalized to the cytosol in a portion of the *pex32* cells (Fig. 3A), while cytosolic fluorescence was occasionally observed in *pex23* and *pex24* cells, but not in *pex29* cells. In *pex32* cultures, typically three types of cells could be discriminated, namely i) cells with a GFP spot without cytosolic fluorescence, ii) cells with a GFP spot in conjunction with cytosolic GFP and iii) cells with only cytosolic GFP.

**Figure 3.**
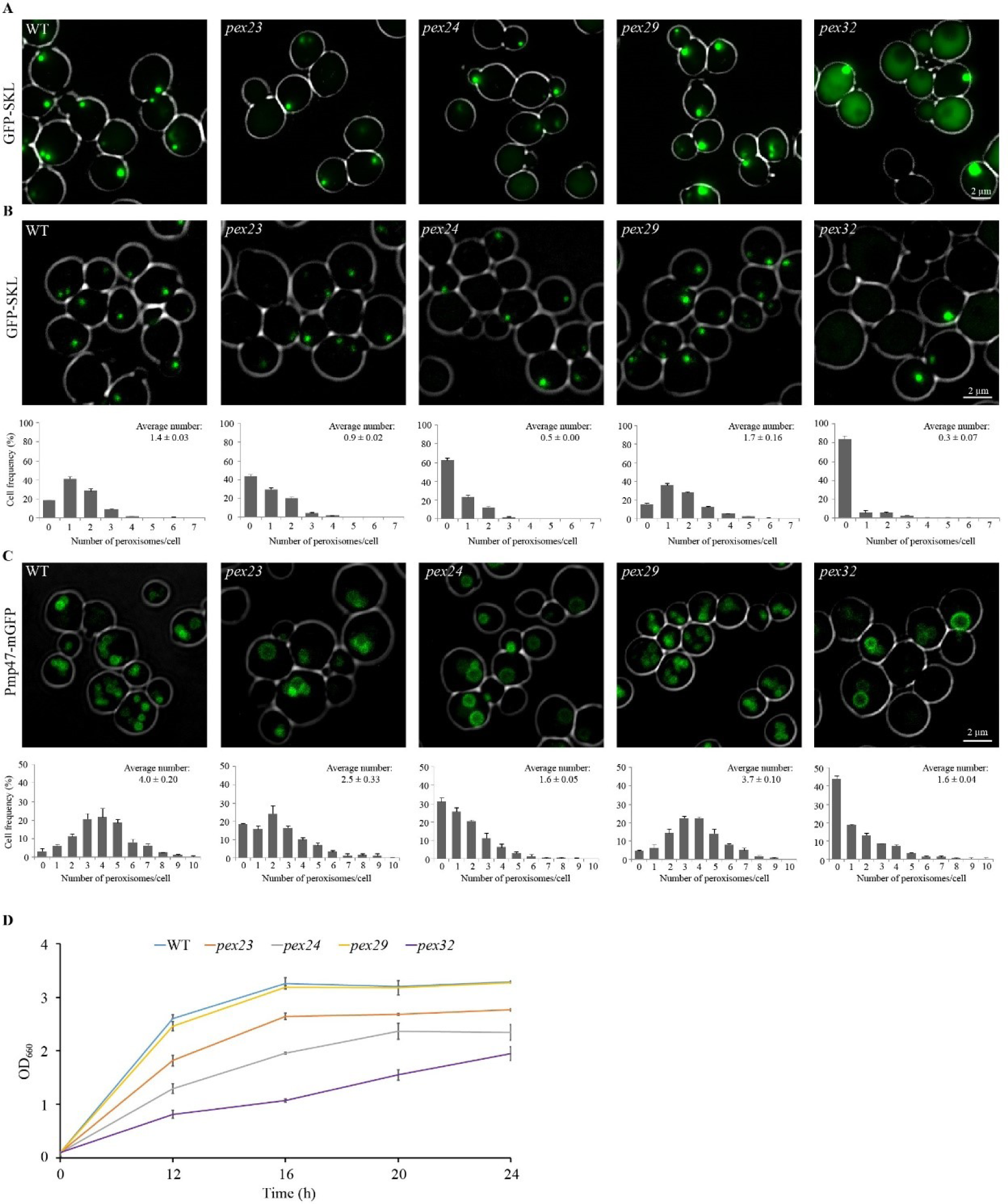
Deletion of *PEX23, PEX24* or *PEX32* results in aberrant peroxisome formation. FM **(A)** and CLSM **(B**) images in conjunction with peroxisome quantification of the indicated deletion strains producing the peroxisome matrix protein GFP-SKL and grown on glucose. 2×500 cells from two independent cultures were quantified. The error bars represent SD from two independent cultures. **(C)** CLSM images of Pmp47-GFP producing cells grown on a mixture of glycerol and methanol. 2×300 cells from two independent cultures were quantified. In the upper right corners of the graphs, the average number of peroxisomes per cell is indicated. The error bars represent SD from two independent cultures. **(D)** Growth curves of the indicated strains in media containing a mixture of glycerol and methanol. The optical density (Y-axis) is expressed as absorbance at 660 nm (OD_660_). The error bars represent the standard deviation (SD) of two independent experiments.

Next, we quantified the number of GFP containing spots by confocal laser scanning microscopy (CLSM) and a custom-made plugin for ImageJ (Thomas et al., 2015). In these images cytosolic fluorescence was not detected in any of the strains due to the lower sensitivity of CLSM relative to wide field FM. The average number of GFP spots per cell was similar in WT and *pex29* cells but reduced in the other three deletion strains. The strongest reduction was observed in *pex24* and *pex32* cells (Fig. 3B). Frequency distribution graphs show that these reductions are accompanied by an increase in the percentage of cells lacking a GFP spot (Fig. 3B).

Finally, we analyzed the strains at peroxisome inducing growth conditions (methanol). Mislocalization of peroxisomal matrix enzymes affects methylotrophic growth (van der Klei et al., 2006). We therefore routinely grow peroxisome-deficient mutants on a mixture of glycerol and methanol (Knoops et al., 2014). At these conditions, cells grow on glycerol (which does not require peroxisome functions), while methanol is used as additional carbon and energy source, depending on the severity of the peroxisome function defect. Growth experiments using glycerol/methanol media revealed the strongest growth defects for the *pex32* and *pex24* strains, while *pex23* showed a minor growth defect and *pex29* grew like WT (Fig. 3D). Quantification of structures marked with the peroxisomal membrane marker Pmp47-GFP indicated that also at these growth conditions peroxisome abundance was reduced, especially in *pex24* and *pex32* cells (Fig. 3C). Moreover, CLSM revealed that *pex23, pex24* and *pex32* cells frequently contained a peroxisome of enhanced size (Fig. 3C).

In conclusion, *pex24* and *pex32* cells showed the most severe peroxisomal phenotypes, while *pex29* cells were like WT and *pex23* cells had minor peroxisomal defects.

### The absence of Pex23 family proteins disrupts peroxisome-ER contacts

In *S. cerevisiae*, Pex23 family proteins and Inp1 play a role in the formation of ER-peroxisome contacts (David et al., 2013; Knoblach et al., 2013; Mast et al., 2016). Using electron microscopy (EM) we analyzed the role of *H. polymorpha* Pex23 proteins and Inp1 in the formation of these contacts (Fig. 4). In WT controls the distance between the peroxisomal and ER membranes was less than 10 nm in approximately 80 % of the peroxisomal profiles, indicating that these represent contact sites (Fig. 4A). In cells of the *pex23* and *pex29* strains this percentage decreased to approximately 40 %, while it further dropped in *pex24* and *pex32* cells to 10 - 20 % (Fig. 4A, B). These changes were not related to a decrease in total cortical ER, which instead slightly increased (Fig. 4D).

**Figure 4.**
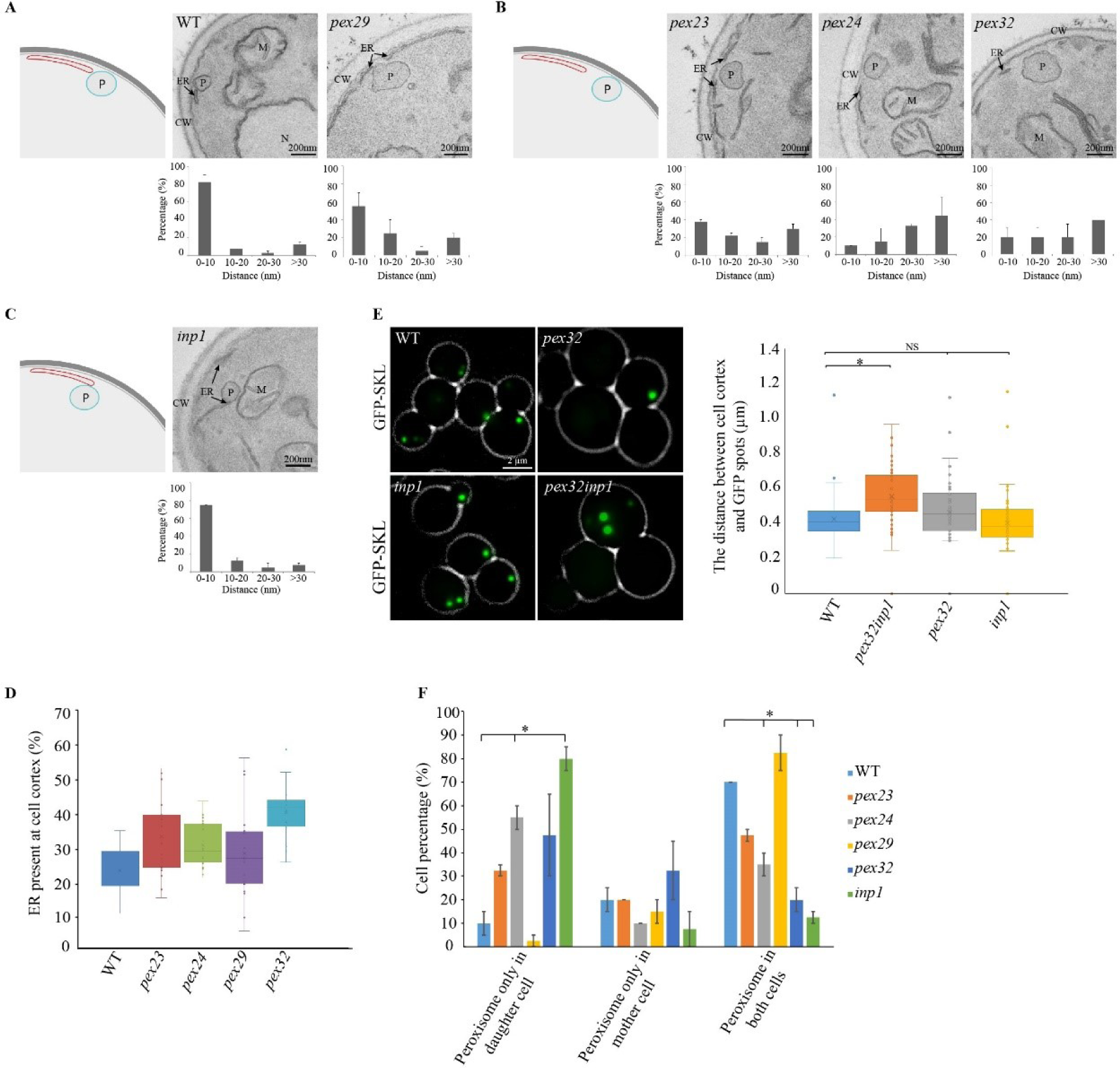
Deletion of *PEX23, PEX24* or *PEX32* results in an increase in distance between peroxisomal and ER membranes. **(A, B, C)** EM images of thin sections of KMnO_4_-fixed glucose-grown cells of the indicated strains and quantification of the distance between the ER and peroxisomal membranes. The error bar represents the SD. 2×21 peroxisomes in random sections from two independent cultures were analyzed. CW - cell wall; ER - endoplasmic reticulum; P - peroxisome; M - mitochondrion; N - nucleus **(D)** Quantification of ER abundance at the cell cortex. The percentage of the cell cortex covered by the ER measured in 20 random cell sections using EM. **(E)** FM images and quantification of the distance between the cell cortex and the GFP spots in the indicated deletion strains producing GFP-SKL and grown on glucose. 2×24 cells from two independent cultures were quantified. **(F)** Quantification of the presence of peroxisomes in both the mother cell and bud in the indicated mutants. The error bar represents the SD. 2×20 peroxisome contained budding yeast cells from two independent cultures were quantified.

Deletion of *INP1* had no effect on the distance at ER-peroxisome contact sites (Fig. 4C), in line with our recent observation that *H. polymorpha* Inp1 associates peroxisomes to the plasma membrane (Wu, 2020).

FM analysis of the position of peroxisomes demonstrated that peroxisomes remained close to the cell cortex upon deletion of either *PEX32* or *INP1*. However, in a *pex32 inp1* double mutant peroxisomes were more frequently observed in the central part of the cells, indicating that Pex32 and Inp1 both contribute to the cortical association of peroxisomes (Fig. 4E).

Analysis of peroxisome segregation in budding cells showed that in the *pex29* strain peroxisomes segregated like in the WT control. However, in *pex24* cultures a large fraction of the budding cells contained peroxisomes solely in the buds (Fig. 4F). In addition, in *pex24* and *pex32* cultures the percentage of budding cells with a peroxisome in both the mother cell and bud strongly reduced relative to the WT control (Fig. 4F).

These data show that close associations between peroxisomes and the ER require Pex23 family proteins, of which Pex24 and Pex32 are paramount. Together with Inp1, Pex23 family proteins contribute to peroxisome positioning at the cell cortex and proper segregation in budding cells.

### An artificial ER-peroxisome tether suppresses the peroxisomal phenotypes

To study whether the effect of the absence of Pex24 and Pex32 on peroxisome biology is due to the disrupted peroxisome-ER contacts, we introduced an artificial tether in an attempt to re-associate both organelles. This approach is based on studies in *S. cerevisiae*, in which the absence of proteins of the ER-Mitochondria Encounter Structure (ERMES) is partially complemented by artificially anchoring mitochondria to the ER (Kornmann et al., 2009). To this end we constructed an artificial tether protein consisting of full length Pex14 and the tail anchor of the ER protein Ubc6, separated by two heme-agglutinin tags (HA). This construct (P_*ADH1*_Pex14-HA-HA-Ubc6^TA^), termed ER-PER, was introduced in WT and the four deletion strains (Fig. 5A). EM showed that introduction of ER-PER resulted in regions of close opposition (< 10 nm) between the ER/nuclear envelope and the peroxisomal membranes (Fig. 5BC). Immuno-EM using anti-HA antibodies confirmed the presence of ER-PER tether protein at these regions (Fig. 5B). EM also showed that in all strains producing ER-PER multiple peroxisomes were present as in WT controls producing ER-PER upon growth on a mixture of methanol and glycerol (Fig. 5C), which was confirmed by FM (Fig. 5D). Also, in *pex32* cells with the ER-PER and producing GFP-SKL, cytosolic fluorescence was not detectable (Fig. 5D). Peroxisome quantification showed that peroxisome numbers in *pex24* and *pex32* cells containing ER-PER were similar as in WT control cells producing ER-PER (Fig. 5E; compare Fig. 3C).

**Figure 5.**
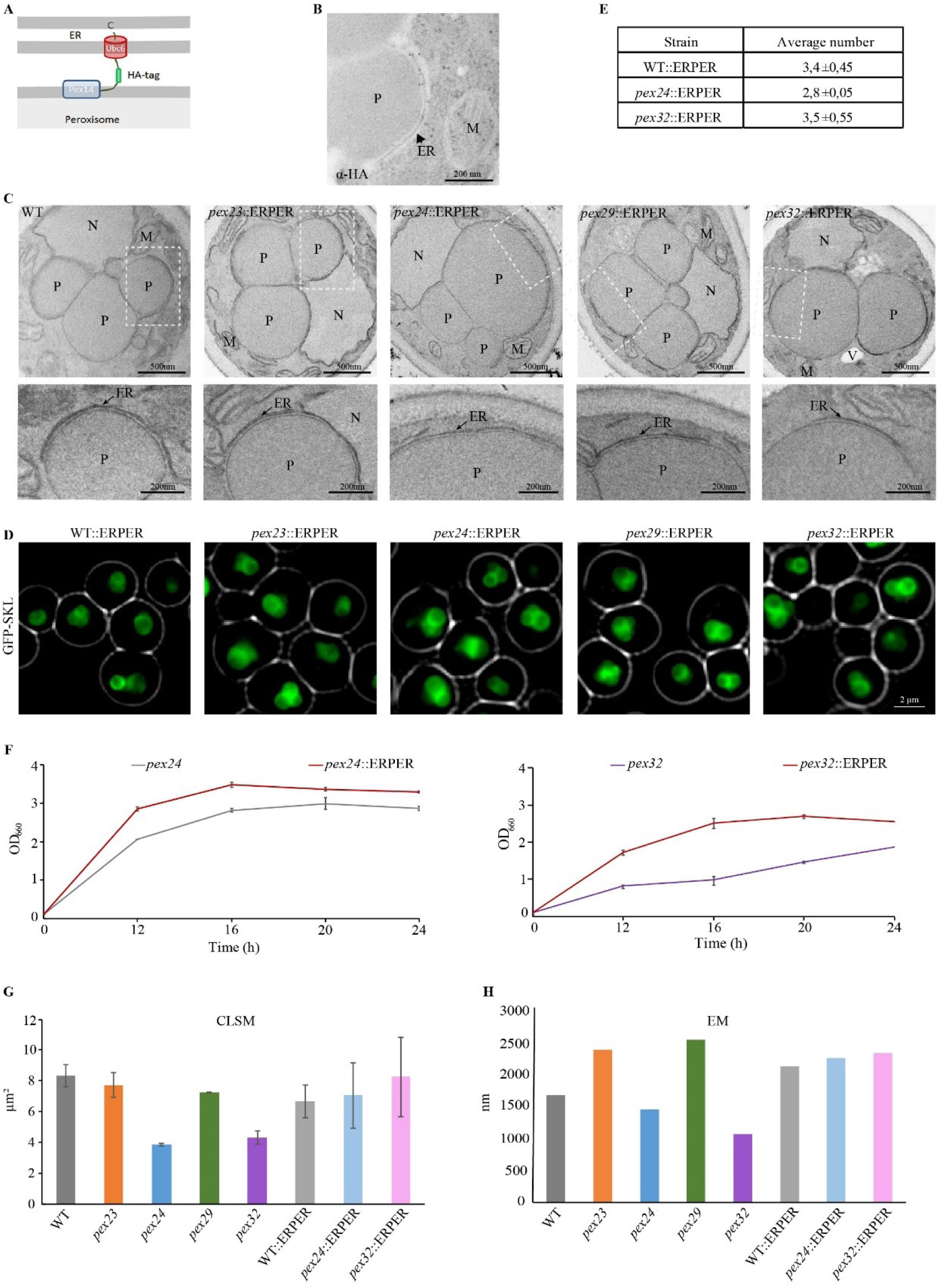
Suppression of peroxisome defects by an artificial ER-peroxisome tether. **(A)** Schematic representation of the ER-PER tethering. **(B)** Immunolabelling using HA antibodies of a WT P_ADH1_*PEX14*-*HAHA*-*UBC6* cell. M - mitochondrion; P - peroxisome; ER-endoplasmic reticulum. **(C)** EM images of KMnO_4_-fixed cells of the indicated mutants strains producing ER-PER. **(D)** Fluorescence microscopy analysis of the indicated strains grown on glycerol/methanol and producing GFP-SKL. Growth curves of the indicated strains. Error bars indicate SD of two independent experiments. (**E**) Quantification of peroxisome numbers based on CLSM analysis of methanol/glycerol grown PMP47-GFP producing cells of the indicated mutant strains containing ER-PER. 2×300 cells from two independent cultures were quantified. (**F**) Growth curves of the indicated strains on glycerol/methanol. (**G**) Average cellular peroxisome surface area calculation based on CLSM images of methanol/glycerol-grown cells of the indicated strains. 2×300 cells have been quantified. Error bars indicate SD of two independent experiments. (**H**) Quantification of the average abundance of peroxisomal membranes in 50 cell sections of the indicated strains from single experiment.

Introduction of ER-PER partially suppressed the growth defects that were observed for the *pex24* and *pex32* deletion strains on glycerol/methanol media (Fig. 5F, compare Fig, 3D), indicating that the tether restored peroxisome function. The tether did not alter the capacity of WT and *pex29* cells to grow on methanol (Suppl. Fig. S1). Also, the minor growth defect of *pex23* was not suppressed by ER-PER (Suppl. Fig. S1).

Introduction of P_*ADH1*_Pex14, which does not tether peroxisomes to the ER, did not alter peroxisome biogenesis or function in WT cells (Suppl. Fig. S1). Only introduction of ER-PER (P_*ADH1*_Pex14-HA-HA-Ubc6^TA^), but not P_*ADH1*_Pex14, suppressed the growth defect of *pex32* cells on glycerol/methanol, confirming that artificial tethering and not solely the enhanced Pex14 levels are responsible for suppression of the phenotype (Suppl. Fig. S1).

From this we conclude that the severe peroxisome defects in *pex24* and *pex32* cells are related to a loss in tight peroxisome-ER contacts.

### Pex24 and Pex32 are important for peroxisomal membrane growth

ER contact sites have been implicated in lipid transfer. To test whether the Pex24/Pex32 dependent ER-peroxisome contacts are important for expansion of peroxisomal membranes, we compared the average peroxisomal membrane surface per cell in the four deletion strains relative to the WT control. The plug-in for the analysis of CLSM images allows quantifying the average diameter of peroxisomes by fitting spheres in data obtained from the green channel of combined z-slices of glycerol/methanol grown, PMP47-GFP producing cells (Thomas et al., 2015). From these data we estimated the average peroxisomal membrane surface per cell. As shown in Fig. 5G, these values were reduced in *pex24* and *pex32* cells relative to *pex23, pex29* and WT cells. Because we are aware of the drawbacks of analyzing organelle sizes by FM (the limited resolution of FM may cause an overestimation of the diameter of very small organelles that are more abundant in WT cells), we also quantified the average length of peroxisomal membranes in cell sections using EM (Fig. 5H). This analysis confirmed that in especially in *pex32* cells, but also in *pex24* cells, the peroxisomal membrane surface is reduced.

Similar analyses of the *pex24* and *pex32* strains containing ER-PER showed that the average peroxisome membrane surface area per cell increased again (Fig. 6 GH), suggesting that these proteins may contribute to lipid supply and hence peroxisomal membrane expansion.

**Figure 6.**
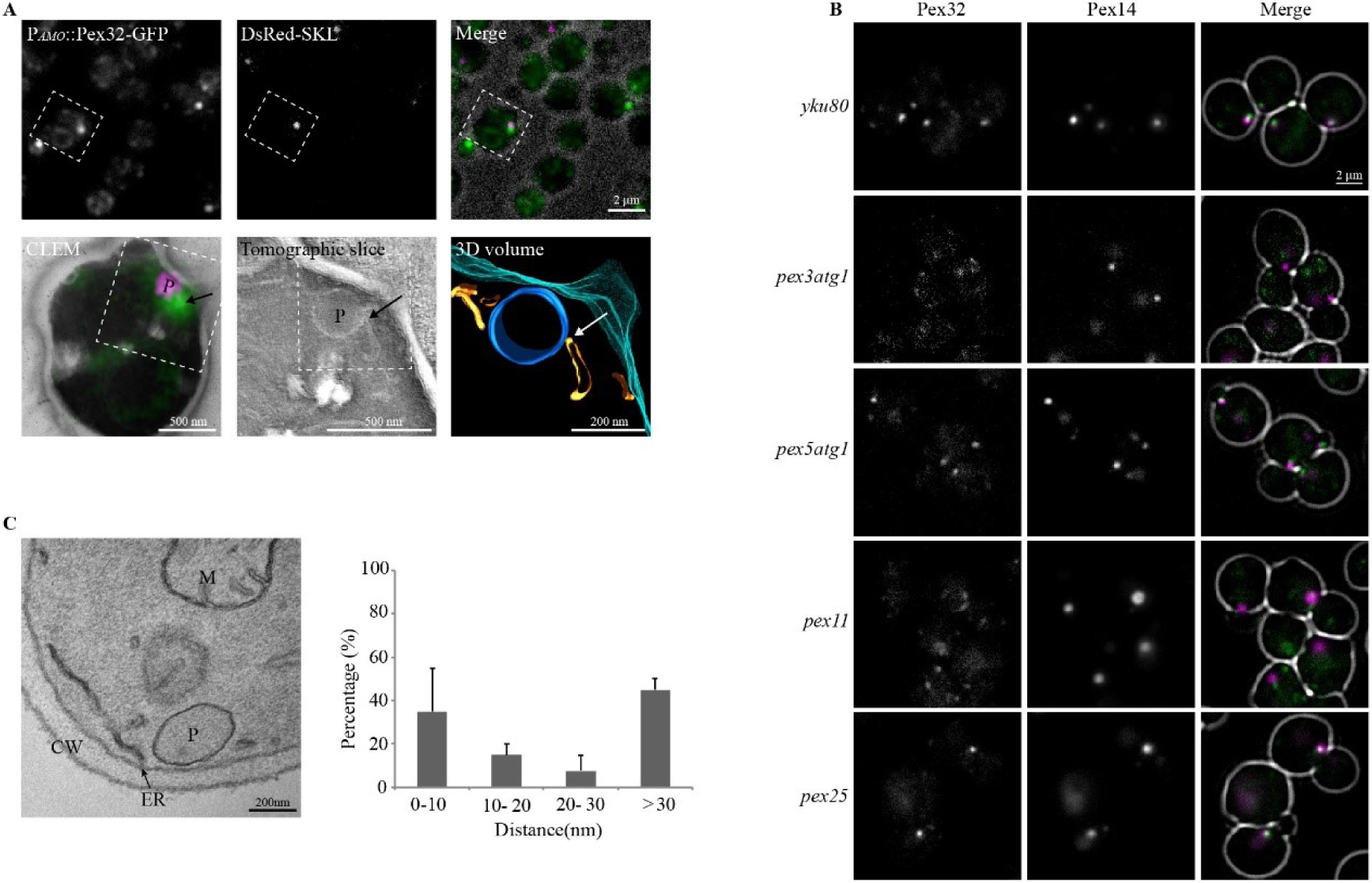
The specific Pex32 localization needs peroxisome membrane protein Pex3 and Pex11. **(A)** CLEM of glucose/methylamine-grown cells producing the peroxisomal matrix marker DsRed-SKL and Pex32-GFP under control of the P_*AMO*_. The upper row shows FM images of 150 nm thick cryo-sections. The lower row shows an overlay of FM and electron microscopy (EM) images of the same cell section. The region of interest is indicated (dashed box). A tomogram was reconstructed and 3D rendered. P – peroxisome: blue; ER: orange; plasma membrane: magenta. **(B)** FM images of glucose-grown indicated *pex* mutant cells producing Pex32-GFP under control of their endogenous promoters together with the peroxisomal marker Pex14-mKate2. **(C)** EM image of KMnO4-fixed glucose-grown *pex11* mutant cell (left) and the ER-peroxisome distance quantification in *PEX11* deletion strain (right). CW - cell wall; ER - endoplasmic reticulum; P - peroxisome; M - mitochondrion. The error bar represents the SD. 2×20 cells from two independent cultures were analyzed.

### The peroxisome membrane protein Pex11 is important for contact site formation

Because *pex32* cells showed the strongest peroxisome phenotype, we studied Pex32 in more detail. Correlative light and electron microscopy (CLEM) was performed to analyze Pex32-GFP localization at high resolution. In order to obtain sufficient fluorescence signal, Pex32-GFP was slightly overexpressed by placing the gene under control of the amine oxidase promoter (P_*AMO*_) and induce this promoter for a short period. At these conditions generally only a single GFP spot was detected per cell. EM analysis revealed that the GFP spot characteristically localizes at the region where the ER and peroxisomal membrane were closely associated (Fig. 6A). In total four tomograms were analyzed and in all the Pex32 spot was present at the ER-peroxisome contact.

Next, we examined whether the peroxisome-ER association is involved in concentrating Pex32 at the contact site. To address this, we localized Pex32-GFP in a *pex3 atg1* double deletion strain, which lacks normal peroxisomes but contains preperoxisomal vesicles (PPVs) (Knoops et al., 2014). In these cells Pex32-GFP accumulation in a spot was lost. Instead, multiple fainter Pex32-GFP spots were observed that showed a typical ER pattern (Fig. 6B). One or a few Pex32-GFP spots were still present in a *pex5 atg1* control strain. In *pex5 atg1* cells small peroxisomes occur that are defective in PTS1 protein import but harbor the complete set of PMPs. Because PPVs in *pex3 atg1* cells and peroxisomes in *pex5 atg1* cells differ in PMP composition, we argued that those PMPs that are absent in PPVs may contribute to the accumulation of Pex32-GFP in spots. One of these is the abundant peroxisomal membrane protein Pex11 (Knoops et al., 2014). FM indicated that in *pex11* cells, but not in *pex25* controls, the bright Pex32-GFP spots were lost (Fig. 6B). Pex25 is also a PMP and belongs to the same protein family as Pex11. Western blot analysis showed that Pex32-GFP levels in these mutants are similar to WT controls, indicating that the absence of the clear Pex32-GFP spots was not due to reduced protein levels (Suppl. Fig. S2).

*H. polymorpha pex11* cells have several features in common with *pex32* cells with respect to growth on methanol, organelle size, number and segregation. Also, these cells typically contain enlarged peroxisomes (Krikken et al., 2009). This led us to examine whether *pex11* cells are have reduced peroxisome-ER contacts. Indeed, EM analysis showed that the distance between ER and peroxisomal membranes increased in *pex11* cells, like in *pex32* cells (Fig. 7C). These data indicate that ER-localized Pex32 together with peroxisomal Pex11 contribute to the formation of peroxisome-ER contacts.

**Figure 7.**
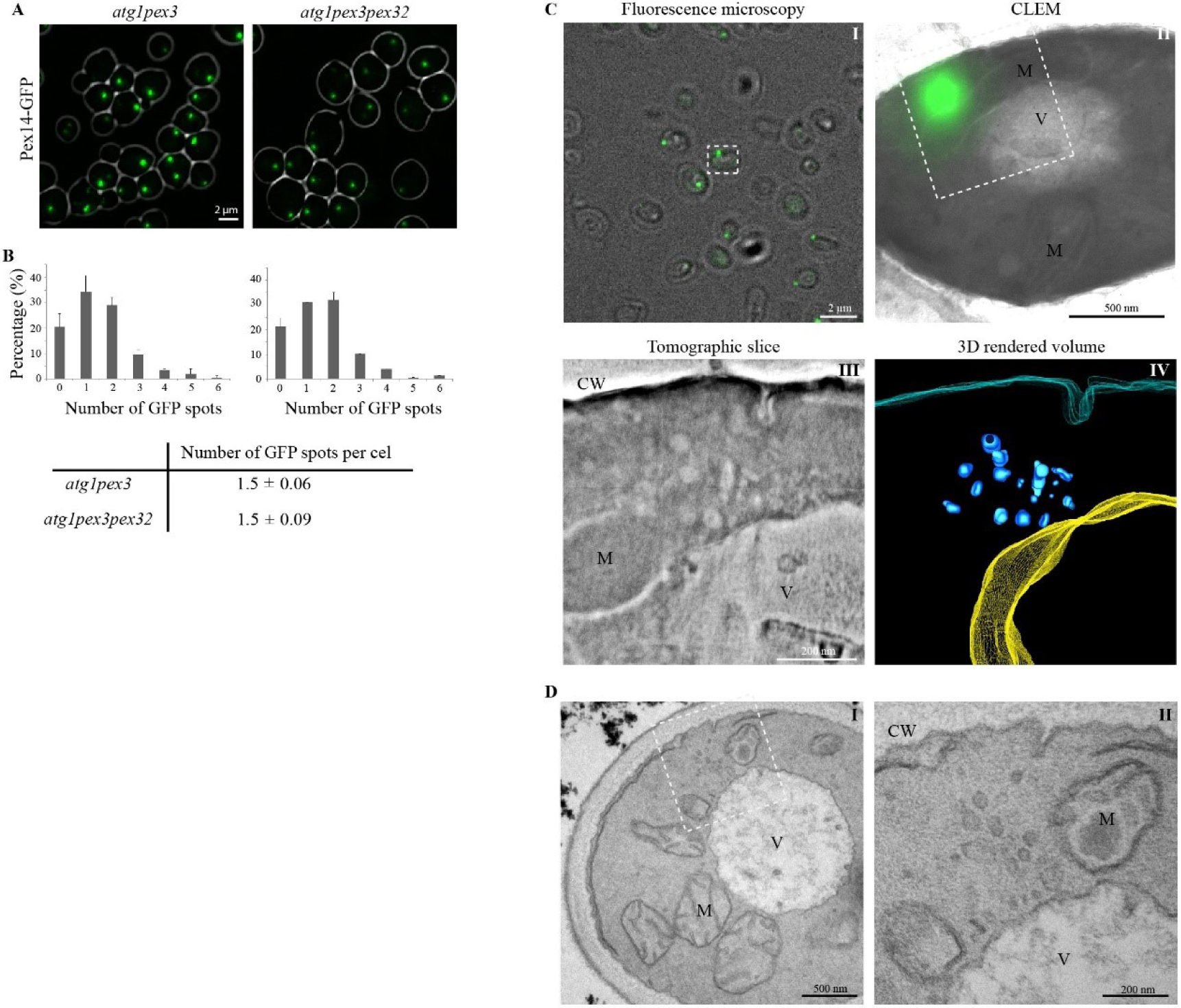
Deletion of *pex32* has no effect on PPV formation. **(A)** CLSM images of glycerol/methanol grown cells producing Pex14-GFP as a PPV marker. **(B)** Distribution of the number of Pex14-GFP spots per cell and the average number of spots per cell quantified from 2x 280 cells per strain. **(C-I)** Wide-field FM image of a thick cryo-section (250 nm) of *atg1 pex3 pex32* cells producing Pex14-GFP. **(C-II)** CLEM showing an overlay image of Pex14-GFP (FM) and the TEM micrograph of the same region indicated with the white dashed box in C-I. **(C-III)** Electron tomographic slice from a tomogram recorded at the region indicated in C-II. White arrows indicate the position of the PPVs. **(C-IV)** 3D rendered volume of the reconstructed tomogram. *Blue*-PPVs, *yellow*- vacuole, *magenta* - plasma membrane. **(D)** EM analysis of KMnO_4_-fixed *atg1 pex3 pex32* cells. D-II shows a higher magnification of the region indicated in D-I.

### The absence of Pex32 does not reduce PPV abundance

In *S. cerevisiae* Pex23 family proteins have been implicated in the formation of PPVs from the ER (Joshi et al., 2016, 2018; Mast et al., 2016). Detailed FM and EM analysis revealed that in *H. polymorpha pex3 atg1* cells deletion of *PEX32* did not result in alterations in PPV abundance (Fig. 7A,B) or morphology (Fig. 7C,D). Therefore, Pex32 is not crucial for PPV formation in *H. polymorpha*.

## Discussion

Here, we report the first systematic study in which all members of the Pex23 protein family of one yeast species are analysed. All four members of the *H. polymorpha* Pex23 protein family (Pex23, Pex24, Pex29 and Pex32) localize to the ER. Of these, Pex32 and Pex24 predominantly accumulate at peroxisome-ER contacts and appeared to be very important for multiple peroxisome features. Pex23 is less important, while we could not detect a peroxisomal phenotype in cells lacking Pex29. Possibly, Pex23 and Pex29 play redundant roles in peroxisome biology or are involved in other functions and hence do not represent true peroxins. Pex23 also accumulates at NVJs, suggesting that Pex23 family proteins may be intrinsic contact site proteins. Initial studies revealed that in *H. polymorpha pex23* and *pex29* cells, but not in *pex24* and *pex32* cells, mitochondrial morphology and lipid body abundance is altered, suggesting that these proteins may contribute to the formation of other organelles (Fei Wu, unpublished observations). Indeed, in *S. cerevisiae* ER domains enriched in Pex30 are the sites where most nascent lipid droplets form (Joshi et al., 2018).

Analysis of an evolutionary tree revealed that HpPex23 proteins can be partitioned in two major subgroups, one containing HpPex23 and HpPex32 and the other HpPex24 and HpPex29. There is no clear correlation between subgroup and molecular function, because the strongest peroxisomal phenotypes occurred in the absence of HpPex24 and HpPex32.

The absence of *H. polymorpha* Pex24 and Pex32 resulted in the loss of peroxisome ER contacts, accompanied by several peroxisome defects. These phenotypes could be suppressed by an artificial peroxisome ER tether protein, indicating that Pex24 and Pex32 function as tethers. The peroxisomal membrane protein Pex11 also contributes to these contacts. Interestingly, *P pastoris* Pex11 pull down experiments resulted in the identification of Pex31, a member of the *P. pastoris* Pex23 protein family (Yan et al., 2008). Moreover, David and colleagues (David et al., 2013) identified ScPex11 as a specific binding partner in ScPex29 complexes, supporting the role of Pex11 in ER-peroxisome contacts. *S. cerevisiae* Pex11 is also a component of a peroxisome-mitochondrion contact site, indicating that Pex11 contributes to the formation of different membrane contacts (Mattiazzi Ušaj et al., 2015).

The loss of peroxisome-ER contacts causes multiple phenotypes. It is not unprecedented that a contact site resident protein is involved in various processes. For instance the vacuolar membrane protein Vac8 functions in NVJs, vacuole fusion and inheritance in *S. cerevisiae* (Pan and Goldfarb, 1998). Also, the mitochondrial outer membrane protein Mdm10 is a component of ERMES and required for membrane protein insertion (Kornmann et al., 2009; Meisinger et al., 2004; Wiedemann and Pfanner, 2017).

A possible function of the Pex24-, Pex32- and Pex11-dependent peroxisome-ER contacts includes transfer of lipids from the ER to peroxisomes. Indeed, we observed reduced peroxisomal membrane surfaces in cells lacking Pex24 or Pex32. Yeast peroxisomes lack lipid biosynthetic enzymes, hence expansion of the peroxisomal membrane relies on the supply of lipids from other sources. In *S. cerevisiae* peroxisomal membrane lipids may originate from multiple sources, including the mitochondrion, the Golgi apparatus, the vacuole and the ER (Flis et al., 2015; Rosenberger et al., 2009). Evidence for non-vesicular lipid transport to peroxisomes has been reported (Raychaudhuri and Prinz, 2008) as well as vesicular transport (Van der Zand and Tabak, 2013).

In glucose-grown *H. polymorpha* cells the single peroxisome invariably associates with the edge of cortical ER sheets, where the ER is highly curved (Wu et al., 2019). Using CLEM we showed that Pex32 specifically localizes to these regions. This is consistent with studies in *S. cerevisiae*, which revealed that members of the Pex23 family occur in complexes with the ER-shaping reticulons, Rtn1/Rtn2, and Yop1(David et al., 2013; Joshi et al., 2016; Mast et al., 2016). ER-shaping proteins have been implicated in lipid exchange between the ER and mitochondria in *S. cerevisiae* (Voss et al., 2012). Therefore, it is tempting to speculate that highly curved ER regions where *H. polymorpha* Pex32 localizes functions in lipid transport. Also, like *S. cerevisiae* Pex30 and Pex31 HpPex32 has a reticulon-like domain and thus may have membrane shaping properties (Joshi et al., 2016). Peroxisome-ER contact sites that contribute to phospholipid transport recently have been identified in mammals as well. At these sites, the ER proteins VAPA/B interact with the peroxisome membrane proteins ACBD4/5 (Costello et al., 2017a, 2017b; Hua et al., 2017).

Another role of the ER contacts may be in peroxisome fission. Mitochondrion-ER contacts are important in the selection of fission sites (Friedman et al., 2011). A comparable mechanism may occur for peroxisomes. This is suggested by presence of enlarged peroxisomes in *pex24* and *pex32* as well as in *pex11* cells, known to be defective in peroxisome fission (Williams et al., 2015). A possible alternative explanation for the presence of the enlarged peroxisomes in *H. polymorpha pex23, pex24* and *pex32* cells is a change in membrane lipid composition, which may interfere with peroxisome fission. Although the absence of *S. cerevisiae* Pex30 changed the ER phospholipid composition (Wang et al., 2018), it is yet unknown whether this peroxin influences the phospholipid content of the peroxisomal membrane.

The ER-peroxisome contacts described in this study also contribute to peroxisome positioning at the cell cortex and proper segregation of the organelles over mother cells and buds. So far, only yeast Inp1 was implicated in peroxisome retention (Fagarasanu et al., 2005; Krikken et al., 2009). We here show that HpPex24 contributes to peroxisome retention as well. HpPex11 is important for peroxisome retention as well, underscoring its role in the formation of peroxisome-ER contacts (Krikken et al., 2009).

Earlier studies revealed that the absence of two Pex23 family members in *S. cerevisiae*, Pex30 or Pex31, results in enhanced *de novo* peroxisome formation (David et al., 2013; Mast et al., 2016). This *de novo* pathway has been proposed to occur in yeast *pex3* mutants upon reintroduction of the *PEX3* gene. We recently showed that PPVs are present in *H. polymorpha* and *S. cerevisiae pex3* strains upon blocking autophagy by deletion of *ATG1* (Knoops et al., 2014; Wróblewska et al., 2017). These structures still contain a subset of PMPs, including Pex14. Studies by Joshi et al. (Joshi et al., 2016) showed that deletion of *PEX30* or *PEX31* resulted in a significant decrease in the number of Pex14-GFP spots in *S. cerevisiae pex3 atg1* cells. However, in *H. polymorpha* deletion of *PEX32* in *pex3 atg1* cells affect the abundance of PPVs.

In conclusion, our data indicate that Pex23 family proteins are contact sites tethers. In *H. polymorpha* Pex24 and Pex32 are most important for tethering peroxisomes to the ER, important for several organelle related properties.

## Materials and methods

### Strains and growth conditions

The *H. polymorpha* strains used in this study are listed in Table S1. Yeast cells were grown in batch cultures at 37°C on mineral media (MM) (Van Dijken et al., 1976) supplemented with 0.5% glucose or 0.5% methanol or a mixture of 0.5% methanol and 0.05% glycerol (MM-M/G) as carbon sources and 0.25% ammonium sulfate or 0.25% methylamine as nitrogen sources. When required, amino acids were added to the media to a final concentration of 30 µg/mL. Transformants were selected on YND plates (0.67% yeast nitrogen base without amino acids (YNB; Difco; BD) and 0.5% glucose) or on YPD plates (1% yeast extract, 1% peptone and 1% glucose) containing 2% agar supplemented with 100 µg/mL zeocin (Invitrogen), 300 µg/mL hygromycin B (Invitrogen) or 100 µg/ml nourseothricin (WERNER BioAgents).

### Construction of *H. polymorpha* strains

The plasmids and primers used in this study are listed in Tables S2 and S3. All plasmid integrations were performed as described previously (Faber et al., 1994). All integrations were confirmed by PCR and all deletions were confirmed by PCR and southern blotting.

### Construction of strains expressing Pex23-mGFP, Pex24-mGFP, Pex29-mGFP and Pex32-mGFP under control of the endogenous promoter

A plasmid encoding Pex23-mGFP was constructed as follows: a PCR fragment encoding the C-terminus of *PEX23* was obtained using primers Pex23 GFP-fw and Pex23 GFP-rev with *H. polymorpha* NCYC495 genomic DNA as a template. The obtained PCR fragment was digested with *Bgl*II and *Hin*dIII, and inserted between the *Bgl*II and *Hin*dIII sites of plasmid pHIPZ-mGFP fusinator. *Bsm*BI-linearized pHIPZ *PEX23*-mGFP was transformed into *yku80* cells, producing strain Pex23-mGFP.

The same methods were used to construct Pex24-mGFP, Pex29-mGFP and Pex32-mGFP strains. PCR was performed on WT genomic DNA with primers Pex24 fw and Pex24 rev to get C-terminus of *PEX24*, primers Pex29 fw and Pex29 rev were used for PCR to get C-terminus of *PEX29*, primers Pex32 fw and Pex32 rev were used for PCR to get C-terminus of *PEX32*. The obtained PCR fragment of *PEX24* was digested with *Bgl*II and *Hin*dIII, the PCR fragment of *PEX29* and the PCR fragment of *PEX32* were restricted by *Bam*HI and *Hin*dIII, these three digested fragments were inserted between the *Bgl*II and *Hin*dIII sites of pHIPZ-mGFP fusinator plasmid, respectively. *Bcl*I-linearized pHIPZ *PEX24*-mGFP, *Nru*I-linearized pHIPZ *PEX29*-mGFP and *Mfe*I-linearized pHIPZ *PEX32*-mGFP were transformed into *yku80* cells separately, producing strains Pex24-mGFP, Pex29-mGFP and Pex32-mGFP. *Mun*I-linearized pHIPH *PEX14*-mKate2 was transformed into Pex23-mGFP, Pex24-mGFP, Pex29-mGFP and Pex32-mGFP cells for colocalization study.

For Pex23 family proteins colocalization study with the ER: *Dra*I-linearized pHIPX7 BiP_*N30*_-mCherry-HDEL was integrated into Pex24-mGFP and Pex29-mGFP cells, respectively. *Stu*I-linearized pHIPX7 BiP_*N30*_-mCherry-HDEL was transformed into Pex23-mGFP cells and Pex32-mGFP cells, respectively. Plasmid pHIPX7 BiP_*N30*_-mCherry-HDEL was constructed as follows: first, a PCR fragment containing *BiP* was obtained with primers KN18 and KN19 using WT genomic DNA as templates. The obtained fragment was digested with *Bam*HI and *Hin*dIII, inserted between the *Bam*HI and *Hin*dIII sites of pBlueScript II, resulting in plasmid pBS-BiP. Then a PCR fragment containing GFP-HDEL was obtained with primers KN14 and KN17 using pANL29 as templates, the resulting fragment was digested with *Sal*I and *Bgl*II, and then inserted between the *Sal*I and *Bgl*II sites of pBS-BiP, resulting in pBS-BiP_*N30*_-GFP-HDEL. Subsequently, pBS-BiP_*N30*_-GFP-HDEL was digested with *Bam*HI/*Sal*I and inserted between the *Bam*HI/*Sal*I sites of pHIPX7 to obtain pHIPX7 BiP_*N30*_-GFP-HDEL. Plasmid pHIPX7 BiP_*N30*_-GFP-HDEL was digested with *Bam*HI/*Eco*RI and inserted between the *Bam*HI/*Eco*RI sites of pHIPX4, resulting in pHIPX4 BiP_*N30*_-GFP-HDEL. *Not*I and *Sal*I were used to digest pHIPX4 BiP_*N30*_-GFP-HDEL and inserted between the *Not*I and *Sal*I sites of pHIPZ4 DsRed-SKL to obtain plasmid pRSA017. Later, a PCR fragment was obtained by primers BIPmCh1_fw and BIPmCh1_rev on plasmid pMCE02, the resulting fragment was inserted between *Bgl*II and *Sal*I sites of pRSA017 to obtain pHIPZ4 BiP_*N30*_-mCherry-HDEL. Finally, a PCR fragment was obtained by primers BIPmCh2_fw and BIPmCh1_rev using plasmid pHIPZ4 BiP_*N30*_-mCherry-HDEL as a template, the resulting fragment was inserted between *Bgl*II and *Sal*I sites of pHIPX7 BiP_*N30*_-GFP-HDEL, resulting in pHIPX7 BiP_*N30*_-mCherry-HDEL.

### Construction of strains producing Pex23-mGFP, Pex24-mGFP, Pex29-mGFP and Pex32-mGFP under control of the P_*AMO*_

A plasmid encoding Pex24-mGFP behind the strong inducible promoter amine oxidase was constructed as follows: a PCR fragment containing *PEX24*-mGFP was obtained using primers Pex24GFP fw and Pex24GFP rev with Pex24-mGFP genomic DNA as template. This PCR product and pHIPH5 were restricted by *Sbf*I and *Bam*HI and ligated which resulted in pHIPH5 *PEX24*-mGFP. *Pml*I-linearized pHIPH5 *PEX24*-mGFP was transformed into *yku80* cells to produce strain P_*AMO*_Pex24-mGFP. Plasmid pHIPH5 was constructed by *Not*I and *Sph*I digested pHIPZ5, inserted into the *Not*I and *Sph*I sites of pHIPH4.

The plasmid pHIPH5 *PEX29*-mGFP and plasmid pHIPH5 *PEX32*-mGFP were constructed in the same way. Primers Pex29ov-fw and Pex29ov-rev were used to get a PCR fragment containing *PEX29*-mGFP with Pex29-mGFP genomic DNA as the template. Primers Pex32ov-fw and Pex32ov-rev were used to get a PCR fragment containing *PEX32*-mGFP with Pex32-mGFP genomic DNA as the template. PCR products of *PEX29*-mGFP and *PEX32*-mGFP were restricted by *Sbf*I and *Bcl*I, and insert between the *Sbf*I and *Bcl*I sites of pHIPH5 *PEX24*-mGFP, respectively, to get plasmid pHIPH5 *PEX29*-mGFP and pHIPH5 *PEX32*-mGFP. *Nar*I-linearized pHIPH5 *PEX29*-mGFP and pHIPH5 *PEX32*-mGFP were integrated into *yku80* cells separately to produce strain P_*AMO*_Pex29-mGFP and P_*AMO*_Pex32-mGFP.

The plasmid of pHIPH5 *PEX23*-mGFP was constructed in two steps: first, a PCR fragment containing partial (no start codon) *PEX23*-mGFP was obtained using primers Pex23ov-fw and Pex23ov-rev with Pex23-mGFP genomic DNA as a template. PCR product and pHIPH5 *PEX24*-mGFP were restricted by *Sbf*I and *Bam*HI, ligated to produce pHIPH5 *PEX23p*-mGFP. Next, a PCR using primers Pex23ov2-fw and Pex23ov2-rev to obtain the left partial (with start codon) *PEX23*-mGFP fragment with plasmid pHIPH5 *PEX24*-mGFP as templates. PCR product and pHIPH5 *PEX23p*-mGFP were restricted by *Not*I and *Bam*HI, ligated to produce pHIPH5 *PEX23*-mGFP. *Nar*I-linearized pHIPH5 *PEX23*-mGFP was transformed into *yku80* cells to produce P_*AMO*_Pex23-mGFP.

*Eco*RI-linearized pHIPN18 DsRed-SKL was integrated into *yku80*, P_*AMO*_Pex23-mGFP, P_*AMO*_Pex24-mGFP, P_*AMO*_Pex29-mGFP, P_*AMO*_Pex32-mGFP cells, respectively. A plasmid encoding pHIPN18 DsRed-SKL was constructed as follows: a vector fragment was obtained by *Hin*dIII and *Sal*I digestion of pHIPN18 GFP-SKL, whereas the DsRed-SKL insertion fragment was obtained by *Hin*dIII and *Sal*I digestion of pHIPZ4 DsRed-SKL, ligation resulted in the plasmid pHIPN18 DsRed-SKL. Plasmid pHIPN18 GFP-SKL was constructed as follow: *Not*I and *Xba*I digested pAMK94 inserted into the *Not*I and *Xba*I sites of pHIPN4 to get pHIPN18 GFP-SKL. Plasmid pAMK94 was constructed as follow: a PCR fragment containing *ADH1* was amplified with primers ADH1 fw and ADH1 rev with WT genomic DNA as template. *Not*I and *Hin*dIII digested PCR product was inserted into *Not*I and *Hin*dIII sites of pHIPZ4 eGFP-SKL.

*Mun*I-linearized pHIPN *VAC8*-mKate2 was integrated into Pex23-mGFP and P_*AMO*_Pex24GFP cells to produce Vac8-mKate2. Plasmid pHIPN *VAC8*-mKate2 was constructed by fragment ligation from *Hin*dIII/*Sal*I digested plasmid pHIPZ *VAC8*-mKate2 and *Hin*dIII/*Sal*I digested plasmid pHIPN *PEX14*-mCherry. Plasmid pHIPZ *VAC8*-GFP and plasmid pHIPZ *PEX14*-mKate2 were digested with *Hin*dIII and *Bgl*II and ligated to obtain plasmid pHIPZ *VAC8*-mKate2. Plasmid pHIPZ *VAC8*-GFP was constructed by amplification of the *VAC8* gene, lacking the stop codon, using primers Vac8_BglII R and Vac8_F and genomic DNA as template. The resulting PCR product was digested with *Hin*dIII and *Bgl*II, and ligated between the *Hin*dIII and *Bgl*II sites of the pHIPZ-mGFP fusinator plasmid.

### Construction of *pex23, pex24, pex29* and *pex32* deletion strains

The *pex23* deletion strain was constructed by replacing the *PEX23* region with the zeocin resistance gene as follows: first, a PCR fragment containing the zeocin resistance gene and 50 bp of the *PEX23* flanking regions were amplified with primers PEX23-Fw and PEX23-Rev using plasmid pENTR221-zeocin as template. The resulting *PEX23* deletion cassette was transformed into *yku80* cells to obtain strain *pex23. PEX24, PEX29* and *PEX32* were also replaced by the zeocin resistance gene in the same way. Primers for *PEX24* deletion cassette were PEX24-Fw and PEX24-Rev, primers for *PEX29* deletion cassette were dPEX29-F and dPEX29-R, and primers for *PEX32* deletion cassette were dPEX32-F and dPEX32-R. These three deletion cassettes were transformed into y*ku80* cells, respectively, producing *pex24, pex29* and *pex32*.

The *Stu*I-linearized pHIPN7 GFP-SKL was transformed into *pex23* and *pex24* mutant cells separately to produce GFP-SKL. The *Ahd*I-linearized pFEM35 was transformed into *yku80, pex29* and *pex32* mutant cells, respectively, producing GFP-SKL.

The *Mun*I-linearized pHIPN *PMP47*-mGFP plasmid was transformed into *pex23, pex24, pex29* and *pex32* cells, respectively. Plasmid pHIPN *PMP47*-mGFP was constructed as follows: a PCR fragment encoding the nourseothricin resistance gene was obtained with primers Nat-fwd and Nat-rev using plasmid pHIPN4 as a template. The obtained PCR fragment was digested with *Not*I and *Xho*I and inserted between the *Not*I and *Xho*I sites of pMCE7, resulting in plasmid pHIPN *PMP47*-mGFP.

### Construction of *pex23* family mutants with or without an artificial ER-PER tether

To introduce an artificial peroxisome-ER tether, two plasmids pARM115 (pHIPH18 *PEX14*) and pARM118 (pHIPH18 *PEX14*-2HA-*UBC6*) were constructed as follows. A PCR fragment containing *PEX14* was amplified with primers Pex14-HindIII-fw and Pex14-PspXI-rev using the WT genomic DNA as a template. The PCR fragment was digested with *Hin*dIII and *Psp*XI, inserted between the *Hin*dIII and *Sal*I sites of pAMK94 to get plasmid pHIPZ18 *PEX14*. A *Not*I/*Bpi*I digested fragment from plasmid pHIPZ18 *PEX14* and a *Not*I/*Bpi*I digested fragment from plasmid pHIPH4 were ligated, resulting in plasmid pARM115. The *Age*I-linearized was transformed into *yku80*∷GFP-SKL and *pex32*∷GFP-SKL cells to produce P_*ADH1*_Pex14 (Pex14++). A PCR fragment containing *PEX14*-2xHA was amplified by primers HindIII-Pex14 and Pex14-HA-HA. A fragment containing 2xHA-*UBC6* was amplified with primers HAHA-Ubc6 and Ubc6-PspXI and WT genomic DNA as template. The obtained PCR fragments were purified and used as templates together with primers HindIII-Pex14 and Ubc6-PspXI in a second PCR reaction. The obtained overlap PCR fragment was digested with *Hin*dIII and *Psp*XI, and inserted between the *Hin*dIII and *Sal*I sites of pAMK94, resulting in plasmid pARM053 (pHIPZ18 *PEX14*-2HA-*UBC6*). A *Not*I/*Bpi*I digested fragment from plasmid pAMK053 and a *Not*I/*Bpi*I digested fragment from plasmid pHIPH4 were ligated, resulting in plasmid pARM118. Then the *Age*I-linearized pARM118 was transformed into *yku80*∷GFP-SKL, *yku80*∷Pmp47-GFP, *pex23*∷GFP-SKL, *pex24*∷GFP-SKL, *pex24*∷Pmp47-GFP, *pex29*∷GFP-SKL, *pex32*∷GFP-SKL and *pex32*∷Pmp47-GFP cells, respectively, to produce P_*ADH1*_Pex14-2HA-Ubc6 (ERPER).

### Expression of Pex32-mGFP in different *pex* mutant cells

The *Bgl*II-linearized pHIPZ *PEX32*-mGFP were transformed into *pex3 atg1*∷Pex14-mCherry, *pex5 atg1*∷Pex14-mCherry, *pex11* and *pex25* cells, respectively, to produce Pex32-mGFP. *Blp*I-linearized pARM014 (pHIPX7 *PEX14*-mCherry) was transformed into *pex5 atg1* cells, which resulted in *pex5 atg1*∷Pex14-mCherry. Plasmid pARM014 was constructed with following steps: first, a PCR fragment containing Pex14-mCherry was amplified with primers PRARM001 and PRARM002 using pSEM01 as a template. The obtained PCR fragment was digested with *Not*I and *Hin*dIII, and inserted between the *Not*I and *Hin*dIII sites of plasmid pHIPX7, resulting in plasmid pARM014. *ATG1* deletion cassette was amplified by PCR with primers pDEL-ATG1-fwd + pDEL-ATG1-rev and plasmid pARM011 as template. Then the PCR product integrated into *pex5* to get *pex5 atg1* mutant.

Two plasmids allowing disruption of *H. polymorpha PEX25* were constructed using Multisite Gateway technology as follows: First, the 5’ and 3’ flanking regions of the *PEX25* gene were amplified by PCR with primers RSAPex25-1+RSAPex25-2 and RSAPex25-3+RSAPex25-4, respectively, using *H. polymorpha* NCYC495 genomic DNA as a template. The resulting fragments were then recombined in donor vectors pDONR P4-P1R and pDONR P2R-P3, resulting in plasmids pENTR-PEX25 5’ and pENTR-PEX25 3’, respectively. Then, PCR amplification was performed using primers attB1-Ptef1-forward and attB2-Ttef1-reverse using pHIPN4 as the template. The resulting PCR fragment was recombined into vector pDONR-221 yielding entry vector pENTR-221-NAT. Recombination of the entry vectors pENTR-PEX25 5’, pENTR-221-NAT, and pENTR-PEX25 3’, and the destination vector pDEST-R4-R3, resulted in pRSA018. Then *PEX25* disruption cassette containing neursothricin resistance gene was amplified with primers RSAPex25-5 and RSAPex25-6 using pRSA018 as a template. To create *pex25*, the *PEX25* disruption cassette was transformed into *yku80* cells. *Blp*I-linearized pHIPH *PEX14*-mCherry was integrated into *pex11*∷Pex32-mGFP or *pex25*∷Pex32-mGFP to produce Pex14-mCherry.

### Construction of *pex32 inp1* double and *pex3 atg1 pex32* triple deletion strains

To construct *pex32 inp1* mutant, a PCR fragment containing *INP1* deletion cassette was amplified with primers dInp1FW-F and dInp1-REV using plasmid pHIPH5 as a template. The resulting *INP1* deletion cassette was transformed into *pex32* cells to get double deletion of *pex32 inp1*. The *Ahd*I-linearized pFEM35 was transformed into *pex32 inp1* to produce GFP-SKL.

To construct *pex3 atg1 pex32* strain, a PCR fragment containing *PEX32* deletion cassette was amplified with primers dPex32-F and dPex32-R using *pex32* genomic DNA as a template. The resulting *PEX32* deletion cassette was transformed into *pex3 atg1* cells to get triple mutant of *pex3 atg1 pex32. Xho*I-linearized pHIPN-*PEX14*-mGFP plasmid was integrated into *pex3 atg1 pex32* cells.

A plasmid encoding pHIPN *PEX14*-mGFP was constructed as follows: a PCR fragment containing the nourseothricin resistance gene was obtained using primers Nat fw and Nat rev with plasmid pHIPN4 as a template. The PCR product and pSNA12 were digested with *Nsi*I and *Not*I, then ligated to produce pHIPN *PEX14*-mGFP.

### Molecular and biochemical techniques

DNA restriction enzymes were used as recommended by the suppliers (Thermo Fisher Scientific or New England Biolabs). Polymerase chain reactions (PCR) for cloning were carried out with Phusion High-Fidelity DNA Polymerase (Thermo Fisher Scientific). An initial selection of positive transformants by colony PCR was carried out using Phire polymerase (Thermo Fisher Scientific). For DNA and amino acid sequence analysis, the Clone Manager 5 program (Scientific and Educational Software, Durham, NC) was used. For western blot analysis, 4 hours glucose-grown cell extracts, 4 hours glucose/methylamine-grown cell extracts or 16 hours glycerol/methanol-grown cell extracts were prepared of whole cells as described previously (Baerends et al., 2000). Equal amounts of protein were loaded per lane and blots were probed with rabbit polyclonal antisera against *H. polymorpha* Pex14, or pyruvate carboxylase 1 (Pyc1). GFP fusion proteins of Pex23, Pex24, Pex29, and Pex32 were detected using mouse monoclonal antiserum against GFP (sc-9996; Santa Cruz Biotechnology, Inc.). Secondary goat anti–rabbit or goat anti–mouse antibodies conjugated to horseradish peroxidase (Thermo Fisher Scientific) were used for detection. Pyc1 was used as a loading control. Blots were scanned by using a densitometer (GS-710; Bio-Rad Laboratories).

### Fluorescence microscopy

Wide-field FM images on living cells and on cryosections for CLEM were captured at room temperature using a 100×1.30 NA objective (Carl Zeiss, Oberkochen, Germany). Images were obtained from the cells in growth media using a fluorescence microscope (Axioscope A1; Carl Zeiss), Micro-Manager 1.4 software and a digital camera (Coolsnap HQ^2^; Photometrics). The GFP fluorescence were visualized with a 470/40 nm band-pass excitation filter, a 495 nm dichromatic mirror, and a 525/50 nm band-pass emission filter. DsRed fluorescence were visualized with a 546/12 nm band-pass excitation filter, a 560 nm dichromatic mirror, and a 575-640 nm band-pass emission filter. mCherry and mKate2 fluorescence were visualized with a 587/25 nm band-pass excitation filter, a 605 nm dichromatic mirror, and a 670/70 nm band-pass emission filter.

Confocal images were captured with an LSM800 Airyscan confocal microscope (Carl Zeiss) using Zen 2.3 software (Carl Zeiss) and a 100x/1.40 plan apochromat objective and GaAsP detectors. For quantitative analysis of peroxisomes or Pex14-mGFP fluorescent spots, z-stacks were made of randomly chosen fields.

Image analysis was performed using ImageJ, and figures were prepared using Adobe Illustrator software.

### Electron microscopy

For morphological analysis, cells were fixed in 1.5% potassium permanganate, post-stained with 0.5% uranyl acetate and embedded in Epon. Image analysis and distance measurements are performed using ImageJ. For the quantification of the ER, the total length of the plasma membrane and the peripheral ER was measured from cell sections and from this the percentage of the cortex covered by the ER was calculated. Correlative light and electron microscopy (CLEM) was performed using cryo-sections as described previously (Knoops et al., 2015). After fluorescence imaging, the grid was post-stained and embedded in a mixture of 0.5% uranyl acetate and 0.5% methylcellulose. Acquisition of the double-tilt tomography series was performed manually in a CM12 TEM running at 100 kV and included a tilt range of 40° to −40 with 2.5° increments. To construct the CLEM images, pictures taken with FM and EM were aligned using the eC-CLEM plugin in Icy (Paul-Gilloteaux et al., 2017) (http://icy.bioimageanalysis.org). Reconstruction of the tomograms was performed using the IMOD software package.

Immuno-EM was performed as described previously (Thomas et al., 2018). Labeling of HA was performed using monoclonal antibodies (Sigma-Aldrich H9658) followed by goat-anti-mouse antibodies conjugated to 6 nm gold (Aurion, the Netherlands).

### In silico analyses

Homologous sequences were detected using BLASTP with an e-value of 1e-5 (Altschul et al., 1990). Linear and secondary structure predictions were realized using Foundation (Bordin et al., 2018).

### Phylogenetic tree

The multiple sequence alignment used as input was created using ClustalOmega (Sievers et al., 2011) with default parameters and manually curated in Jalview (Waterhouse et al., 2009). The tree was generated using PhyML 3.1 (Guindon et al., 2010) using the LG matrix, 100 bootstraps, tree and leaves refinement, SPR moves, and amino acids substitution rates determined empirically.

### Peroxisome membrane surface area calculation

For peroxisome membrane surface area calculation: average peroxisome volume (V) and average peroxisome number per cell (N) were recorded by the plugin for ImageJ (Thomas et al., 2015) from two independent experiments (2×300 cells were counted). Formula 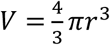 was used to calculate peroxisome radius (r) and formula *S* = 4*πr*^2^ was used to calculate average peroxisome surface area. The average peroxisome number per cell *N* multiply *S* is the amount of peroxisome membrane surface area per cell.

### Quantification of the distance between Pex14-mGFP spots and cell cortex

For the distance between Pex14-mGFP spots and cell cortex quantification, cells containing GFP spots were first selected and processed, then the distance between the middle of the GFP spot and the ring of the cell was measured by using Image J from two independent experiments (2×24 cells were counted). For cells contain two or more GFP spots, only take the GFP spot which nearest to cell ring into account.

### Peroxisome inheritance quantification

Peroxisome inheritance quantification was using the same method as published previously (Krikken et al., 2009). Random pictures of peroxisomal contained budding cells were taken. Assuming yeast cells to be spherical, cells for which the bud volume was less than 25% of the mother cell volume were counted as a budding cell. Quantification experiments were performed using two independent cell cultures (20 cells per culture).

## Author contributions

FW, AA, AMK, RdB and IvdK conceived the project; FW, AA, AMK, RdB, NB, DPD, IJvdK performed the experiments, analysed the data and prepared the figures; FW, IJvdK wrote the original draft. All contributed to reviewing and editing the manuscript.

## Acknowledgements

We thank Malgorzata Krygowska for assistance in strain construction and Tim Levine (University College London, UK) for advice in protein sequence analysis.

## Funding sources and disclosure of conflicts of interest

This work was supported by a grant from the Marie Curie Initial Training Networks (ITN) program PerFuMe (Grant Agreement Number 316723) to NB, DPD and IvdK and the CHINA SCHOLARSHIP COUNCIL (CSC) to FW.

The authors declare no competing financial interests.

