## supplemental figures and tables for "Pex24 and Pex32 tether peroxisomes to the ER for organelle biogenesis, positioning and segregation"

### Figure S1

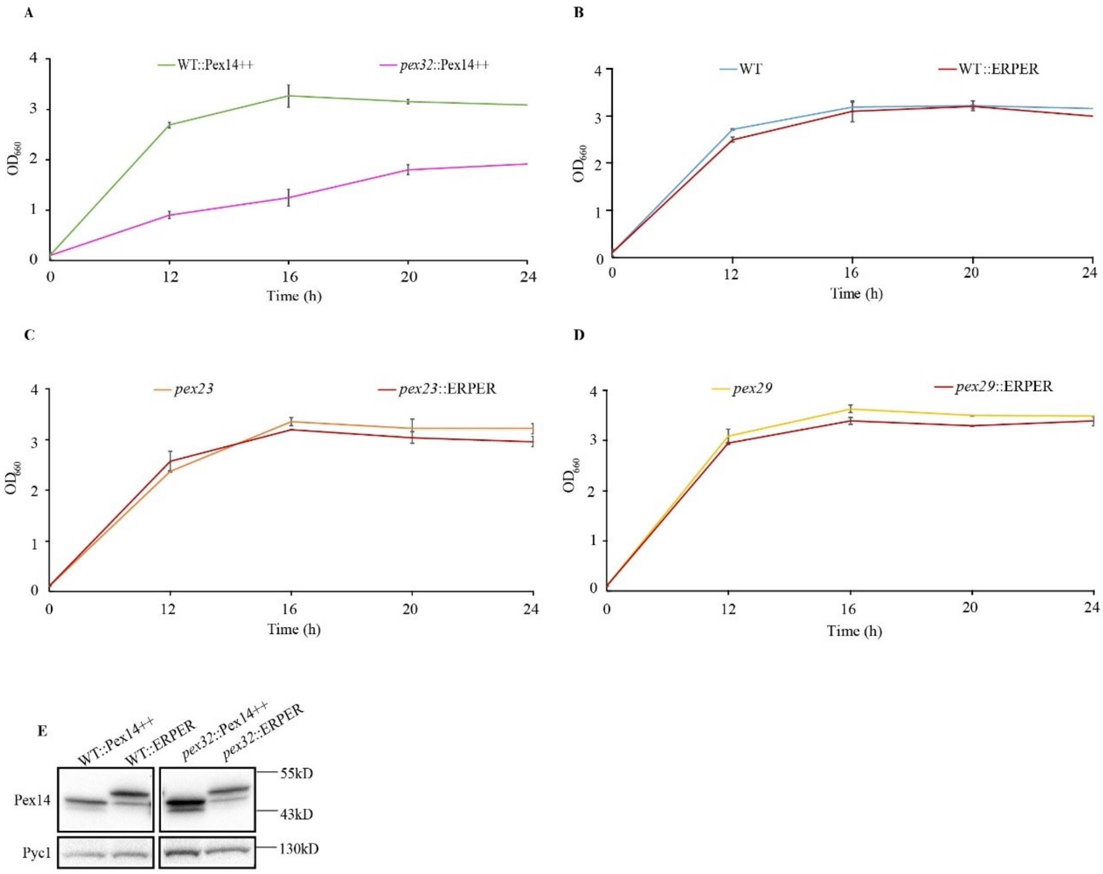


**Figure S1. Growth curve of the indicated strains on glycerol/methanol media. (A)** WT or *pex32* cells overproducing full length Pex14; **(B)** WT control cells with or without ER-PER; **(C)** *pex23* cells with and without ER-PER; **(D)** *pex29* cells with and without ER-PER. Error bars indicate SD from two independent experiments. **(E)** Western blot of WT and *pex32* mutant cells producing $P_{ADH1}$*PEX14* (Pex14++) or ER-PER using α-Pex14 antibodies, showing that similar levels of full length Pex14 or ER-PER were obtained. Pyc1 was used as loading control.

#### Figure S2

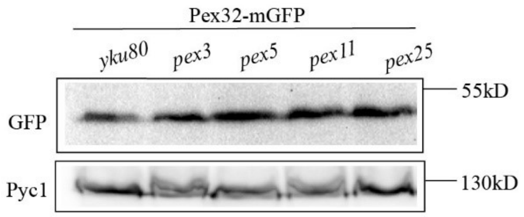


**Figure S2.** Western blot showing Pex32-GFP levels in glucose grown cells of the indicated strains. The blot was decorated with α-GFP antibodies. Pyc1 was used as loading control.

**Supplementary Table 1. Strains used in this study**

| Strain | Characteristics | Reference |
|---|---|---|
| WT | NCYC495; *leu 1.1* | (Sudbery et al., 1988) |
| WT::*yku80* | NCYC495 *yku80* deletion strain; *leu 1.1*, *URA3* | (Saraya et al., 2012) |
| Pex23-mGFP | *yku80* with pHIPZ_*PEX23*-mGFP; *leu 1.1*,*URA3*, $Zeo^R$ | This study |
| Pex24-mGFP | *yku80* with pHIPZ_*PEX24*-mGFP; *leu1.1, URA3*, $Zeo^R$ | This study |
| Pex29-mGFP | *yku80* with pHIPZ_*PEX29*-mGFP; *leu1.1, URA3*, $Zeo^R$ | This study |
| Pex32-mGFP | *yku80* with pHIPZ_*PEX32*-mGFP; *leu1.1, URA3*, $Zeo^R$ | This study |
| Pex23-mGFP::Pex14-mKate2 | Pex23-mGFP with pHIPH_*PEX14*-mKate2; *leu 1.1*, *URA3*, $Zeo^R$, $Hph^R$ | This study |
| Pex24-mGFP::Pex14-mKate2 | Pex24-mGFP with pHIPH_*PEX14*-mKate2; *leu 1.1*, *URA3*, $Zeo^R$, $Hph^R$ | This study |
| Pex29-mGFP::Pex14-mKate2 | Pex29-mGFP with pHIPH_*PEX14*-mKate2; *leu 1.1*, *URA3*, $Zeo^R$, $Hph^R$ | This study |
| Pex32-mGFP::Pex14-mKate2 | Pex32-mGFP with pHIPH_*PEX14*-mKate2; *leu 1.1*, *URA3*, $Zeo^R$, $Hph^R$ | This study |
| Pex24-mGFP::BiP-mCherry-HDEL | Pex24-mGFP with pHIPX7 $BiP_{N30}$-mCherry-HDEL; *URA3*, $Zeo^R$, *LEU2* | This study |
| Pex29-mGFP::BiP-mCherry-HDEL | Pex29-mGFP with pHIPX7 $BiP_{N30}$-mCherry-HDEL; *URA3*, $Zeo^R$, *LEU2* | This study |
| Pex23-mGFP::BiP-mCherry-HDEL | Pex23-mGFP with pHIPX7 $BiP_{N30}$-mCherry-HDEL; *URA3*, $Zeo^R$, *LEU2* | This study |
| Pex32-mGFP::BiP-mCherry-HDEL | Pex32-mGFP with pHIPX7 $BiP_{N30}$-mCherry-HDEL; *URA3*, $Zeo^R$, *LEU2* | This study |
| $P_{AMO}$Pex24-mGFP | *yku80* with pHIPH5 *PEX24*-mGFP; *leu 1.1*, *URA3*, $Hph^R$ | This study |
| $P_{AMO}$Pex29-mGFP | *yku80* with pHIPH5 *PEX29*-mGFP; *leu 1.1*, *URA3*, $Hph^R$ | This study |
| $P_{AMO}$Pex32-mGFP | *yku80* with pHIPH5 *PEX32*-mGFP; *leu 1.1*, *URA3*, $Hph^R$ | This study |
| $P_{AMO}$Pex23-mGFP | *yku80* with pHIPH5 *PEX23*-mGFP; *leu 1.1*, *URA3*, $Hph^R$ | This study |
| WT::$P_{ADH1}$DsRed-SKL | *yku80* with pHIPN18 DsRed-SKL; *leu 1.1*, *URA3*, $Nat^R$ | This study |
| $P_{AMO}$Pex23-mGFP::DsRed-SKL | $P_{AMO}$Pex23-mGFP with pHIPN18 DsRed-SKL; *leu 1.1*, *URA3*, $Hph^R$, $Nat^R$ | This study |
| $P_{AMO}$Pex24-mGFP::DsRed-SKL | $P_{AMO}$Pex24-mGFP with pHIPN18 DsRed-SKL; *leu 1.1*, *URA3*, $Hph^R$, $Nat^R$ | This study |

| | | |
|---|---|---|
| $P_{AMO}$Pex29-mGFP::DsRed-SKL | $P_{AMO}$Pex29-mGFP with pHIPN18 DsRed-SKL; *leu 1.1*, *URA3*, $Hph^R$, $Nat^R$ | This study |
| $P_{AMO}$Pex32-mGFP::DsRed-SKL | $P_{AMO}$Pex32-mGFP with pHIPN18 DsRed-SKL; *leu 1.1*, *URA3*, $Hph^R$, $Nat^R$ | This study |
| Pex23-mGFP::Vac8-mKate2 | Pex23-mGFP with pHIPN_*VAC8*-mKate2; *URA3*, $Zeo^R$, $Nat^R$ | This study |
| $P_{AMO}$Pex24-mGFP::Vac8-mKate2 | $P_{AMO}$Pex24-mGFP with pHIPN_VAC8-mKate2; *leu 1.1*, *URA3*, $Hph^R$, $Nat^R$ | This study |
| *pex23* | *yku80* with *PEX23* deletion strain; *leu 1.1,URA3*, $Zeo^R$ | This study |
| *pex24* | *yku80* with *PEX24* deletion strain; *leu 1.1,URA3*, $Zeo^R$ | This study |
| *pex29* | *yku80* with *PEX29* deletion strain; *leu 1.1,URA3*, $Zeo^R$ | This study |
| *pex32* | *yku80* with *PEX32* deletion strain; *leu 1.1,URA3*, $Zeo^R$ | This study |
| WT::$P_{TEF1}$eGFP-SKL | *yku80* with pFEM35; *URA3*, *LEU2* | (Krikken et al., 2009) |
| *pex23*::$P_{TEF1}$eGFP-SKL | *pex23* with pHIPN7 eGFP-SKL; *leu 1.1*, *URA3*, $Zeo^R$, $Nat^R$, | This study |
| *pex24*::$P_{TEF1}$eGFP-SKL | *pex24* with pHIPN7 eGFP-SKL; *leu 1.1*, *URA3*, $Zeo^R$, $Nat^R$, | This study |
| *pex29*::$P_{TEF1}$eGFP-SKL | *pex29* with pFEM35; *URA3*, $Zeo^R$, *LEU2* | This study |
| *pex32*::$P_{TEF1}$eGFP-SKL | *pex32* with pFEM35; *URA3*, $Zeo^R$, *LEU2* | This study |
| WT::Pmp47-mGFP | *yku80* with pMCE7; *leu 1.1*, *URA3*, $Zeo^R$ | (Manivannan et al., 2013) |
| *pex23*::Pmp47-mGFP | *pex23* with pHIPN_*PMP47*-mGFP; *leu 1.1*, *URA3*, $Zeo^R$, $Nat^R$ | This study |
| *pex24*::Pmp47-mGFP | *pex24* with pHIPN_*PMP47*-mGFP; *leu 1.1*, *URA3*, $Zeo^R$, $Nat^R$ | This study |
| *pex29*::Pmp47-mGFP | *pex29* with pHIPN_*PMP47*-mGFP; *leu 1.1*, *URA3*, $Zeo^R$, $Nat^R$ | This study |
| *pex32*::Pmp47-mGFP | *pex32* with pHIPN_*PMP47*-mGFP; *leu 1.1*, *URA3*, $Zeo^R$, $Nat^R$ | This study |
| WT::$P_{TEF}$GFP-SKL::ERPER | WT::$P_{TEF1}$eGFP-SKL with pHIPH18 *PEX14-2HA-UBC6*; *URA3*, *LEU2*, $Hph^R$ | This study |
| *pex23*::$P_{TEF}$GFP-SKL::ERPER | *pex23*::$P_{TEF1}$eGFP-SKL with pHIPH18 *PEX14-2HA-UBC6*; *leu 1.1*, *URA3*, $Zeo^R$, $Nat^R$, $Hph^R$ | This study |
| *pex24*::$P_{TEF}$GFP-SKL::ERPER | *pex24*::$P_{TEF1}$eGFP-SKL with pHIPH18 *PEX14-2HA-UBC6*; *leu 1.1*, *URA3*, $Zeo^R$, $Nat^R$, $Hph^R$ | This study |
| *pex29*::$P_{TEF}$GFP-SKL::ERPER | *pex29*::$P_{TEF1}$eGFP-SKL with pHIPH18 *PEX14-2HA-UBC6*; *URA3,* $Zeo^R$, *LEU2*, $Hph^R$ | This study |

| | | |
|---|---|---|
| *pex32*::P$_{TEF}$GFP-SKL::ERPER | *pex32*::P$_{TEF1}$eGFP-SKL with pHIPH18 *PEX14-2HA-UBC6*; *URA3,* Zeo$^R$, *LEU2*, Hph$^R$ | This study |
| WT::P$_{TEF}$GFP-SKL::P$_{ADH1}$Pex14 | WT::P$_{TEF1}$eGFP-SKL with pHIPH18 *PEX14*; *URA3*, *LEU2*, Hph$^R$ | This study |
| *pex32*::P$_{TEF}$GFP-SKL::P$_{ADH1}$Pex14 | *pex32*::P$_{TEF1}$eGFP-SKL with pHIPH18 *PEX14*; *URA3,* Zeo$^R$, *LEU2*, Hph$^R$ | This study |
| WT::Pmp47-mGFP::ERPER | WT::Pmp47-mGFP with pHIPH18 *PEX14-2HA-UBC6*; *leu 1.1*, *URA3*, Zeo$^R$, Hph$^R$ | This study |
| *pex24*::Pmp47-mGFP::ERPER | *pex24*::Pmp47-mGFP with pHIPH18 *PEX14-2HA-UBC6*; *leu 1.1*, *URA3*, Zeo$^R$, Nat$^R$, Hph$^R$ | This study |
| *pex32*::Pmp47-mGFP::ERPER | *pex32*::Pmp47-mGFP with pHIPH18 *PEX14-2HA-UBC6*; *leu 1.1*, *URA3*, Zeo$^R$, Nat$^R$, Hph$^R$ | This study |
| *pex3 atg1*::Pex14-mCherry::Pex32-mGFP | *pex3 atg1::*Pex14-mCherry with pHIPZ_*PEX32*-mGFP; *URA3*, *LEU2*, Nat$^R$, Zeo$^R$ | This study |
| *pex5* | *PEX5* deletion strain; *leu 1.1*, URA3 | (van der Klei et al., 1995) |
| *pex5 atg1* | *pex5* with *ATG1* deletion cassette; *leu 1.1*, *URA3*, Hph$^R$ | This study |
| *pex5 atg1*::Pex14-mCherry | *pex5 atg1* with pHIPX_*PEX14*-mCherry; *URA3*, Hph$^R$, *LEU2* | This study |
| *pex5 atg1*::Pex14-mCherry::Pex32-mGFP | *pex5 atg1*::Pex14-mCherry with pHIPZ_*PEX32*-mGFP; *URA3*, *LEU2*, Hph$^R$, Zeo$^R$ | This study |
| *pex11* | *PEX11* deletion strain; *leu 1.1*, *URA3* | (Krikken et al., 2009) |
| *pex11*::Pex32-mGFP | *pex11* with pHIPZ_*PEX32*-mGFP; *leu 1.1*, *URA3,* Zeo$^R$ | This study |
| *pex11*::Pex32-mGFP::Pex14-mCherry | *pex11*::Pex32-mGFP with pSEM01; *leu 1.1*, *URA3,* Zeo$^R$, Hph$^R$ | This study |
| *pex25* | *yku80* with *PEX25* deletion strain; *leu 1.1*, Nat$^R$ | This study |
| *pex25*::Pex32-mGFP | *pex25* with pHIPZ_*PEX32*-mGFP; *leu 1.1*,Zeo$^R$,Nat$^R$ | This study |
| *pex25*::Pex32-mGFP::Pex14-mCherry | *pex25*::Pex32-mGFP with pARM001; *leu 1.1*,Zeo$^R$,Nat$^R$, Hph$^R$ | This study |
| *inp1* | *INP1* deletion strain; *leu 1.1*, *URA3* | (Krikken et al., 2009) |
| *inp1*::GFP-SKL | *inp1* with pHIPX7 GFP-SKL; *URA3*, *LEU2* | (Krikken et al., 2009) |
| *inp1 pex32* | *pex32* with *INP1* deletion cassette; *leu 1.1*, *URA3*, Zeo$^R$, Hph$^R$ | This study |
| *inp1 pex32*::GFP-SKL | *inp1pex32* with pHIPX7 GFP-SKL; *URA3*, Zeo$^R$, *LEU2* | This study |
| *pex3 atg1 pex32* | *yku80* with *PEX3 ATG1 PEX32* triple deletion strain, *URA3*, *LEU2*, Hph$^R$, Zeo$^R$ | This study |
| *pex3 atg1* | *yku80* with *PEX3 ATG1* double deletion strain; | (Knoops et al., 2014) |

| | | |
|---|---|---|
| | *URA3*, *LEU2*, $Hph^R$ | |
| *pex3 atg1*::Pex14-mGFP | *pex3 atg1* with pSNA12; *URA3*, *LEU2*, $Nat^R$ | (Knoops et al., 2014) |
| *pex3 atg1 pex32::*Pex14-mGFP | *pex32 pex3 atg1* with pHIPN_*PEX14*-mGFP; *URA3*, *LEU2*, $Zeo^R$, $Hph^R$, $Nat^R$ | This study |
| *pex3 atg1*::Pex14-mCherry | *pex3 atg1* with pSEM01; *URA3*, *LEU2*, $Nat^R$ | (Knoops et al., 2014) |

**Supplementary Table 2. Plasmids used in this study**

| Plasmid | Characteristics | Reference |
|---|---|---|
| pHIPZ-mGFP fusinator | pHIPZ plasmid containing mGFP and *AMO* terminator; $Amp^R$, $Zeo^R$ | (Saraya et al., 2010) |
| pHIPZ_*PEX23*-mGFP | pHIPZ plasmid containing the C-terminal of *PEX23* fused to mGFP; $Amp^R$, $Zeo^R$ | This study |
| pHIPZ_*PEX24*-mGFP | pHIPZ plasmid containing the C-terminal of *PEX24* fused to mGFP; $Amp^R$, $Zeo^R$ | This study |
| pHIPZ_*PEX29*-mGFP | pHIPZ plasmid containing the C-terminal of *PEX29* fused to mGFP; $Amp^R$, $Zeo^R$ | This study |
| pHIPZ_*PEX32*-mGFP | pHIPZ plasmid containing the C-terminal of *PEX32* fused to mGFP; $Amp^R$, $Zeo^R$ | This study |
| pHIPH_*PEX14*-mKate2 | pHIPH Plasmid containing the C-terminal region of *PEX14* fused to mKate2; $Amp^R$, $Hph^R$ | (Chen et al., 2018) |
| pHIPX7 $BiP_{N30}$-mCherry-HDEL | pHIPX plasmid containing $BiP_{N30}$ fused tomCherry-HDEL under control of *TEF* promoter; $Kan^R$, *LEU2* | This study |
| pBlueScript II | Standard vector; $Amp^R$ | Fermentas |
| pBS-BiP | p-Bluescript II containing *BIP*; $Amp^R$ | This study |
| pANL29 | pHIPZ plasmid containing GFP-SKL under the control of *AOX* promoter; $Amp^R$, $Zeo^R$ | (Leão-Helder et al., 2003) |
| pBS-$BiP_{N30}$-GFP-HDEL | p-Bluescript II containing *BIP*N30- GFP-HDEL; $Amp^R$ | This study |
| pHIPX7 | pHIPX plasmid containing *TEF* promoter; $Kan^R$, *LEU2* | (Baerends et al., 1997) |
| pHIPX7 $BiP_{N30}$-mGFP-HDEL | pHIPX plasmid containing *BIP*N30 fused to GFP-HDEL under the control of *TEF* promoter; $Kan^R$, *LEU2* | This study |
| pHIPX4 | pHIPX plasmid containing *AOX* promoter; $Kan^R$, *LEU2* | (Gietl et al., 1994) |
| pHIPX4 $BiP_{N30}$-GFP-HDEL | pHIPX containing $BIP_{N30}$ fused to GFP-HDEL under the control of *AOX* promoter; $Kan^R$, *LEU2* | This study |
| pHIPZ4 DsRed-SKL | pHIPZ plasmid containing DsRed-SKL under control of *AOX* promoter; $Amp^R$, $Zeo^R$ | (Monastyrska et al., 2005) |
| pRSA017 | pHIPZ containing $BIP_{N30}$ fused to GFP-HDEL under control of *AOX* promoter; $Amp^R$, $Zeo^R$ | This study |
| pMCE02 | pHIPN plasmid containing mCherry; $Amp^R$, $Nat^R$ | (Cepińska et al., 2011) |
| pHIPZ4 $BiP_{N30}$-mCherry-HDEL | pHIPZ containing $BIP_{N30}$ fused to GFP-HDEL under control of *AOX* promoter; $Amp^R$, $Zeo^R$ | This study |
| pHIPH5 | pHIPH plasmid containing *AMO* promoter; $Amp^R$, $Hph^R$ | This study |
| pHIPH5 *PEX24*-mGFP | pHIPH plasmid containing *PEX24* fused with mGFP under the control of *AMO* promotor; $Amp^R$, $Hph^R$ | This study |
| pHIPZ5 | pHIPZ plasmid containing *AMO* promoter; $Amp^R$, $Zeo^R$ | (Faber et al., 1994) |
| pHIPH4 | Plasmid containing *HPH* marker under the control of *AOX* promoter; $Amp^R$,$Hph^R$ | (Saraya et al., 2012) |
| pHIPH5 *PEX29*-mGFP | pHIPH plasmid containing *PEX29*-mGFP under the | This study |

| | | |
|---|---|---|
| | control of *AMO* promotor; $Amp^R$, $Hph^R$ | |
| pHIPH5 *PEX32*-mGFP | pHIPH plasmid containing *PEX32*-mGFP under the control of *AMO* promotor; $Amp^R$, $Hph^R$ | This study |
| pHIPH5 *PEX23p*-mGFP | pHIPH plasmid containing partial (without start code) *PEX23*-mGFP under the control of *AMO* promotor; $Amp^R$, $Hph^R$ | This study |
| pHIPH5 *PEX23*-mGFP | pHIPH plasmid containing *PEX23*-mGFP under the control of *AMO* promotor; $Amp^R$, $Hph^R$ | This study |
| pHIPN18 DsRed-SKL | pHIPN plasmid containing DsRed-SKL under control of *ADH1* promoter; $Amp^R$, $Nat^R$ | This study |
| pHIPN18 GFP-SKL | pHIPN plasmid containing GFP-SKL under control of *ADH1* promoter; $Amp^R$, $Nat^R$ | This study |
| pAMK94 | pHIPZ plasmid containing GFP-SKL under control of *ADH1* promoter; $Amp^R$, $Zeo^R$ | This study |
| pHIPN4 | pHIPN plasmid containing *AOX* promoter; $Amp^R$, $Nat^R$ | (Cepińska et al., 2011) |
| pHIPZ4 eGFP-SKL | pHIPZ plasmid containing GFP-SKL under the control of *AOX* promoter; $Amp^R$, $Zeo^R$ | (Leão-Helder et al., 2003) |
| pHIPN_*VAC8*-mKate2 | pHIPN plasmid containing VAC8 fused with mKate2; $Amp^R$, $Nat^R$ | This study |
| pHIPZ_*VAC8*-mKate2 | pHIPZ plasmid containing VAC8 fused with mKate2; $Amp^R$, $Zeo^R$ | This study |
| pSEM01 | pHIPN plasmid containing C terminal part of *PEX14* fused to mCherry; $Amp^R$, $Nat^R$ | (Knoops et al., 2014) |
| pHIPZ_*VAC8*-mGFP | pHIPZ plasmid containing VAC8 fused with GFP; $Amp^R$, $Zeo^R$ | This study |
| pHIPZ_*PEX14*-mKate2 | pHIPZ plasmid containing the C-terminal part of *PEX14* fused to mKate2; $Amp^R$, $Zeo^R$ | (Chen et al., 2018) |
| pENTR221-zeocin | pDONR221 with s*hble* cassette; $Kan^R$, $Zeo^R$ | (Saraya et al., 2012) |
| pHIPN7 GFP-SKL | pHIPN plasmid containing GFP-SKL under the control of *TEF* promoter; $Amp^R$, $Nat^R$ | (Thomas et al., 2015) |
| pFEM35 | pHIPX plasmid containing GFP-SKL under control of *TEF* promoter; $Kan^R$, *LEU2* | (Krikken et al., 2009) |
| pHIPN_*PMP47*-mGFP | pHIPN plasmid containing C-terminal part of *PMP47* fused to mGFP; $Amp^R$, $Nat^R$ | This study |
| pMCE7 | pHIPZ plasmid containing gene encoding C-terminal part of *PMP47* fused to mGFP; $Amp^R$, $Zeo^R$ | (Cepińska et al., 2011) |
| pARM115 | pHIPH plasmid containing *PEX14* under the contol of *ADH1* promoter; $Amp^R$, $Hph^R$ | This study |
| pARM118 | pHIPH plasmid containing *PEX14*-2HA-*UBC6* under the control of *ADH1* promoter; $Amp^R$,$Hph^R$ | This study |
| pHIPZ18 *PEX14* | pHIPZ plasmid containing *PEX14* under the control of *ADH1* promoter; $Amp^R$, $Zeo^R$ | This study |
| pARM053 | pHIPZ plasmid containing *PEX14*-2xHA-*UBC6* under the | This study |

| | | |
|---|---|---|
| | control of *ADH1* promoter; $Amp^R$, $Zeo^R$ | |
| pARM014 | pHIPX plasmid containing C terminal part of *PEX14* fused to mCherry under the control of *TEF1* promoter; $Kan^R$, *LEU2* | This study |
| pARM011 | Plasmid containing the *ATG1* deletion cassette, $Amp^R$ ,$Hph^R$ | (Thomas et al., 2018) |
| pDONR P4-P1R | Multisite Gateway vector; $Kan^R$, $Cm^R$ | Invitrogen |
| pDONR P2R-P3 | Multisite Gateway vector; $Kan^R$, $Cm^R$ | Invitrogen |
| pENTR-PEX25 5’ | pDONR P4-P1R with 5’ flanking region of PEX25; $Kan^R$ | This study |
| pENTR-PEX25 3’ | pDONR P2R-P3 with 3’ flanking region of PEX25; $Kan^R$ | This study |
| pDONR-221 | Multisite gateway donor vector; $Kan^R$, $Cm^R$ | Invitrogen |
| pENTR-221-NAT | pDONR 221 with NAT cassette; $Nat^R$, $Kan^R$ | This study |
| pDEST-R4-R3 | Multisite Gateway donor vector; $Amp^R$, $Cm^R$ | Invitrogen |
| pRSA018 | Plasmid containing *PEX25* deletion cassette; $Zeo^R$, $Amp^R$ | This study |
| pHIPH_*PEX14*-mCherry | pHIPH plasmid containing C terminal part of *PEX14* fused to mCherry; $Amp^R$, $Hph^R$ | (Thomas et al., 2018) |
| pHIPN_*PEX14*-mGFP | pHIPN plasmid containing C-terminal part of *PEX14* fused to mGFP; $Amp^R$, $Nat^R$ | This study |
| pSNA12 | pHIPZ plasmid containing C-terminal part of *PEX14* fused to mGFP; $Amp^R$, $Zeo^R$ | (Cepińska et al., 2011) |

**Supplementary Table 3. Oligonucleotides used in this study**

| Primer | Sequence |
|---|---|
| Pex23 GFP-fw | CCCAAGCTTGGTGACACGAAAGTTGCTTT |
| Pex23 GFP-rev | AGATCTTCCTTCTTTCTTTTTGTCTGTGACACCACC |
| Pex24 fw | CCCAAGCTTGGATGTCTAATGCCCTACC |
| Pex24 rev | GGAAGATCTTCGCTTTTTGGTGGCCTG |
| Pex29 fw | CCCAAGCTTCCGACAAGCACACCATTCTC |
| Pex29 rev | CGCGGATCCTCCGTCCACAGAATCGATCG |
| Pex32 fw | CCCAAGCTTTAGTGGCGTGCACTGTCCTA |
| Pex32 rev | CGCGGATCCGGTGGTTGCGTCGTCCTCGA |
| KN18 | CCCAAGCTTGGATCCATGTTAACTTTCAATAAGTC |
| KN19 | GGGAAGCTTAGATCTAAACTGCTGTGTTGTTAGTGGG |
| KN14 | CCCCTCGAGAACCTGTACTTCCAGTCGAGATCTGTGAGCAAGGGCGAGGAGC |
| KN17 | GGGGTCGACTTACAGCTCGTCGTGAAGCTTGTACAGCTCG |
| BIPmCh1_fw | GGAAGATCTGTGAGCAAGGGCGAGGAGGA |
| BIPmCh1_rev | GACGTCGACTTAGAGTTCATCATGCTTGTACAGCTCGTCCATGCCGCCGG |
| BIPmCh2_fw | CGCGGATCCATGTTAACTTTCAATAAGTCGG |
| Pex24GFP fw | CGGGATCCATGAGCAATTCTCCTCCTTC |
| Pex24GFP rev | GAGCGACCTGCAGGTTACTTGTACAGCTCGTCCA |
| Pex29ov-fw | GGCGTGATCAATGGAGTCTATGGTAAATAAC |
| Pex29ov-rev | CGACCTGCAGGAGTCGACGCGTGCATGCATG |
| Pex32ov-fw | CGCGGATCCATGTCTGAGCCCAATGTTCG |
| Pex32ov-rev | CGACCTGCAGGTTACTTGTACAGCTCGTCCA |
| Pex23ov-fw | CGGGCCTCATAACATATCTCCG |
| Pex23ov-rev | CGACCTGCAGGTTACTTGTACAGCTCGTCCA |
| Pex23ov2-fw | GGGATGTGCTGCAAGGCGATTAAG |
| Pex23ov2-rev | CGCGGATCCGTAGGCATCTGTACGATATGAAGGACAA |
| ADH1 fw | AAGGAAAAAAGCGGCCGCCCCCTGCATTATTAATCACC |
| ADH1 rev | CCCAAGCTTTTTAAATTGATTGATTGATT |
| Vac8_F | TTGCTGTGGACGAGTCCA |
| Vac8_BglII_R | GAAGATCTCTTGATGAGGTCCAAAATTTG |
| PEX23-Fw | CACCTTCTAGCATTAACAGCAACATTTCAGAAGTACAGCCAACAACAGGCTAA TTCCGATCCCCCACACACCATAGCTTC |
| PEX23-Rev | ATCCATCTTCTGCGTCGCGTATACTTGCTGAACGAATCTTCGGTGGACGGGCAA ATTAAAGCCTTCGAGCGTCCC |
| PEX24-Fw | GTGCACCAGGAGTCCCCAGAAATCATTTGTAGAAATAACTTATCAGACAATTC CGATCCCCCACACACCATAGCTTC |
| PEX24-Rev | TGTTTCAGACGGCTTTTCGATGGCCTGGTTCAGGAATCATAGTTGAGCCCGCAA ATTAAAGCCTTCGAGCGTCCC |
| dPEX29-F | GATTGCGTCTGCAGCAAGTTTACAGAAAATAATTTGTCAACTCTTCCCATGGAG TCTAATTCCGATCCCCCACACACCATAGCTTC |
| dPEX29-R | GTCCTGCCTGGTACGAGAACTTGGTCACAAGATCGTAGCACCATTTCTCGTCCT |

| | |
|---|---|
| | CGGCAAATTAAAGCCTTCGAGCGTCCC |
| dPEX32-F | TCGAGCCATTCAGCTATTTTGGGTCCTTATCCAGTTCTGACTATTTCATCTAATTCCGATCCCCCACACACCATAGCTTC |
| dPEX32-R | TTAGCGTCCAGCCATCTCCACCGGCACGTTGCTTGTGTAATCTCTGGGAAGCAAATTAAAGCCTTCGAGCGTCCC |
| Pex14-HindIII-fw | CCCAAGCTTGGGATGTCTCAACAGCCAGCAAC |
| Pex14-PspXI-rev | GACCTCGAGCTTAGGCATTCAGCTGCCACG |
| HindIII-Pex14 | CCCAAGCTTATGTCTCAACAGCCAGCAAC |
| Pex14-HA-HA | TCCTGCATAGTCCGGGACGTCATAGGGATAGCCCGCATAGTCAGGAACATCGTATGGGTAGGCATTCAGCTGCCACGCCG |
| HAHA-Ubc6 | TACCCATACGATGTTCCTGACTATGCGGGCTATCCCTATGACGTCCCGGACTATGCAGGAGAAAACGGATGGGGCATATA |
| Ubc6-PspXI | CGCCTCGAGCCTATCATCTTGATGTACCTCCGG |
| PRARM001 | ATAGCGGCCGCTTGCAGGAAGTCGACGAAAT |
| PRARM002 | CGGAAGCTTTTACTTGTACAGCTCGTCCA |
| pDEL_ATG1_fwd | ACAGGTCGTTGGTGACTTTAC |
| pDEL_ATG1_rev | CTTCTCGTTGCCCGTGACC |
| RSAPex25-1 | GGGGACAACTTTGTATAGAAAAGTTGCAAAGTCTGGATGGAGGCTTCATCTC |
| RSAPex25-2 | GGGGACTGCTTTTTTGTACAAACTTGAGCGTGGCATGCGGTTCATAGAAAC |
| RSAPex25-3 | GGGGACAGCTTTCTTGTACAAAGTGGGAGTCTCTGCTCGCGTACAAGATC |
| RSAPex25-4 | GGGGACAACTTTGTATAATAAAGTTGACTTGGAGCTGCTGTGCTTGTATG |
| attB1-Ptef1 | GGGGACAAGTTTGTACAAAAAAGCAGGCTGATCCCCCACACACCATAGCTTC |
| attB2-Ttef1 | GGGGACCACTTTGTACAAGAAAGCTGGGTGCTCGTTTTCGACACTGGATGG |
| RSAPex25-5 | CTGGATGGAGGCTTCATCTC |
| RSAPex25-6 | GGAGCTGCTGTGCTTGTATG |
| dInp1FW-F | CTGAAATTCCAACACGAGCTCAACAAAAGCGATGCACACAGCCAGGACGACGCTGGCCAGTCTACCAAGCGGCGCGTGCGGGACATAGTGCGACGGTCGTAAGATCCCCCACACACCATAGC |
| dInp1-REV | CTGCCGTCGCCTTCAAAAGACATCATGGTACTGGAATTGATTGTAGACTCGTTTTCGTCTGTCGCTGCCTTCTCCAGCTTGTCGTCTTTGTCCTCCTCGTCATCATCGATGAATTCGAG |
| dPex32-F | CTTACAACCTAACCGGATGC |
| dPex32-R | GCCAGTTTGCGTTTCCTGTC |
| Nat-fwd | CAAAACCTCGAGACTTGCCTTTGAAGGCTCTT |
| Nat-rev | ATAGTTTAGCGGCCGCATCATCGATGAATTCGAGCT |
